# Evidence for glutamate excitotoxicity that occurs before the onset of striatal cell loss and motor symptoms in an ovine Huntington’s Disease model

**DOI:** 10.1101/2023.06.20.545648

**Authors:** Andrew Jiang, Linya You, Renee R. Handley, Victoria Hawkins, Suzanne J. Reid, Jessie C. Jacobsen, Stefano Patassini, Skye R. Rudiger, Clive J. Mclaughlan, Jennifer M. Kelly, Paul J. Verma, C. Simon Bawden, James F. Gusella, Marcy E. MacDonald, Henry J. Waldvogel, Richard L.M. Faull, Klaus Lehnert, Russell G. Snell

**Author notes:** These two authors contributed equally to this work (Andrew Jiang, Linya You).

## Abstract

**Background:** Huntington’s disease (HD) is a neurodegenerative genetic disorder caused by an expansion in the CAG repeat tract of the huntingtin (*HTT*) gene resulting in a triad of behavioural, cognitive, and motor defects. Current knowledge of disease pathogenesis remains incomplete, and no disease course-modifying interventions are in clinical use. We have previously reported the development and characterisation of the *OVT73* transgenic sheep model of HD. *OVT73* captures an early prodromal phase of the disease with an absence of motor symptomatology even at 5-years of age and no detectable striatal cell loss.

**Methods:** To better understand the disease-initiating events we have undertaken a single nuclei transcriptome study of the striatum of an extensively studied cohort of 5-year-old *OVT73* HD sheep and age matched wild-type controls.

**Results:** We have identified transcriptional upregulation of genes encoding N-methyl-D-aspartate (NMDA), α-amino-3-hydroxy-5-methyl-4-isoxazolepropionic acid (AMPA) and kainate receptors in *OVT73* medium spiny neurons, the cell type preferentially lost early in HD. This observation supports the glutamate excitotoxicity hypothesis as an early neurodegeneration cascade-initiating process. Moreover, we also observed the downstream consequences of excitotoxic stress, including a downregulation of transcription of components for the oxidative phosphorylation complexes. We also found that pathways whose activity has been proposed to reduce excitotoxicity, including the CREB family of transcription factors (*CREB1*, *ATF2, ATF4* and *ATF7*) were transcriptionally downregulated.

**Conclusions:** To our knowledge, the *OVT73* model is the first large mammalian HD model that exhibits transcriptomic signatures of an excitotoxic process in the absence of neuronal loss. Our results suggest that glutamate excitotoxicity is a disease-initiating process. Addressing this biochemical defect early may prevent neuronal loss and avoid the more complex secondary consequences precipitated by cell death.

## Background

Huntington’s disease (HD) is a debilitating neurodegenerative genetic disorder caused by an expanded polyglutamine-encoding CAG repeat in the huntingtin gene (*HTT*) [1]. Individuals with 40 or more CAG codons develop the condition with near complete penetrance [2]. Widespread loss of medium spiny neurons in the caudate nucleus and putamen (striatum) contributes to the presenting symptoms that encompass motor, cognitive and behavioural abnormalities. Despite characterisation of the HD defect and advancements in our understanding of disease pathogenesis, no proposed disease-modifying interventions have proved successful in HD clinical trials.

Glutamate excitotoxicity as an initiator of neuronal death has been a longstanding hypothesis and research focus in neurodegenerative disorders [3–6]. In HD, it has been proposed that the loss of the medium spiny neurons of the striatum is due to elevated synaptic glutamate levels from reduced glutamate clearance and increased glutamate release. This excess synaptic glutamate has been proposed to cause an overactivation of ionotropic glutamate receptors including N-methyl-d-aspartate (NMDA) receptors, α-amino-3-hydroxy-5-methyl-4-isoxazolepropionic acid (AMPA) receptors, and kainate receptors, leading to an influx of calcium ions, chronic membrane depolarisation, oxidative stress, and activation of cell death pathways [3–7]. Evidence for ionotropic glutamate receptor-mediated excitotoxicity was gained through the experimental striatal injection of glutamate receptor agonists (glutamic acid, kainic acid, quinolinic acid) in rodents and non-human primates resulting in medium spiny neuronal degeneration and motor dysfunction [8–14]. Further, studies investigating several HD mouse models provided evidence for an association between excessive NMDA receptor signalling and striatal degeneration [15–17].

Our group has previously generated a transgenic ovine model of HD (named *OVT73*) that expresses the full-length human huntingtin (*HTT*) cDNA with a pure CAG repeat length of 69 codons along with a short CAA CAG tract, resulting in a polyglutamine tract of 73 units. The HTT cDNA is under the control of the 1.1kb human sequence immediately upstream of the transgene [18]. The transcript expression level from the transgene in the *OVT73* animals is estimated to be about ∼10%-20% of a single normal allele seen in HdhQ80, YAC128 and rat BACHD HD models [19]. We hypothesized that this moderate expression level might position this model for the investigation of the initiating stages of HD (prodromal phase). Additionally, the germline transmission of the 73-unit glutamine coding repeat was stable over three generations [19] and somatic instability was observed to be minimal (data not shown). We have animals that are over 10 years of age that do not show any overt symptoms and at 6 years of age that do not exhibit any striatal cell loss [19–23]. Several molecular and behavioural changes comparable to early phase human HD were observed in *OVT73*, including the formation of intracellular *HTT* aggregates [19], circadian rhythm abnormalities [22, 23] and changes in brain, plasma and liver metabolites [20, 21, 24, 25].

To examine the transcriptome-wide changes that may underlie many of these observed molecular changes, we have undertaken RNA-seq of single nuclei from the *OVT73* striatum. The tissue utilised for this study was obtained immediately adjacent to that used for our previously reported bulk RNA-seq analysis, taken from an extensively investigated cohort of 5-year-old animals [19–23]. Multidimensional data from these animals are also publicly available in the form of a queryable database [26]. Surprisingly, we have detected transcriptomic signatures for a glutamate excitotoxicity process occurring in the medium spiny neurons of the *OVT73* striatum, denoted by a widespread upregulation of genes encoding ionotropic glutamate receptors (NMDA, AMPA and Kainate receptors). Moreover, we also observed downstream consequences of excitotoxicity, including a downregulation of genes encoding oxidative phosphorylation complexes. Since these changes were identified in a prodromal model of HD, our results provide the first transcriptomic observations of glutamate excitotoxicity in a large mammalian model in the absence of neuronal loss. These findings corroborate other studies in HD mouse models that suggest excitotoxic stress begins early in the disease, before the onset of symptoms [27–29]. Our results emphasise the importance of early intervention, as waiting until later stages of the disease would be far more challenging due to the various downstream consequences of cell loss. These observations position the *OVT73* HD sheep model as a valuable resource for the development and testing of therapeutics effective in the prodromal phase. Moreover, testing of therapeutics may only be feasible in a model like *OVT73* that is not at a disease stage confounded by the cell death cascade.

## Methods

### Ovine samples

The cohort maintenance and tissue sampling has been reported in previous publications [20, 25]. Briefly, animals were maintained in a certified, purpose-made research facility at the South Australian Research and Development Institute (SARDI). Animals were kept in large paddocks as a mixture of wild-type controls and transgenic animals and fed *ad libitum*. Post mortem striatal samples were taken from six 5-year-old *OVT73* (3 females, 3 males) HD sheep as previously described [20, 25] and six age-matched wild-type controls (2 females, 4 males). Tissue collection was performed in accordance with the SARDI/PIRSA Animal Ethics Committee (approval no. 19/02 and 05/12). All experiments performed adhered to the recommendations in the ARRIVE guidelines [30]. In brief, a lethal dose of pentobarbitone sodium solution (Lethabarb, 1 mL/2 kg body weight) was administrated intravenously, the brains were removed from the skulls, dissected into the 5 distinct blocks (Figure 1) and snap frozen initially on dry ice and then in liquid nitrogen. Samples were wrapped in tinfoil and stored at -80°C until further use.

**Figure 1.**
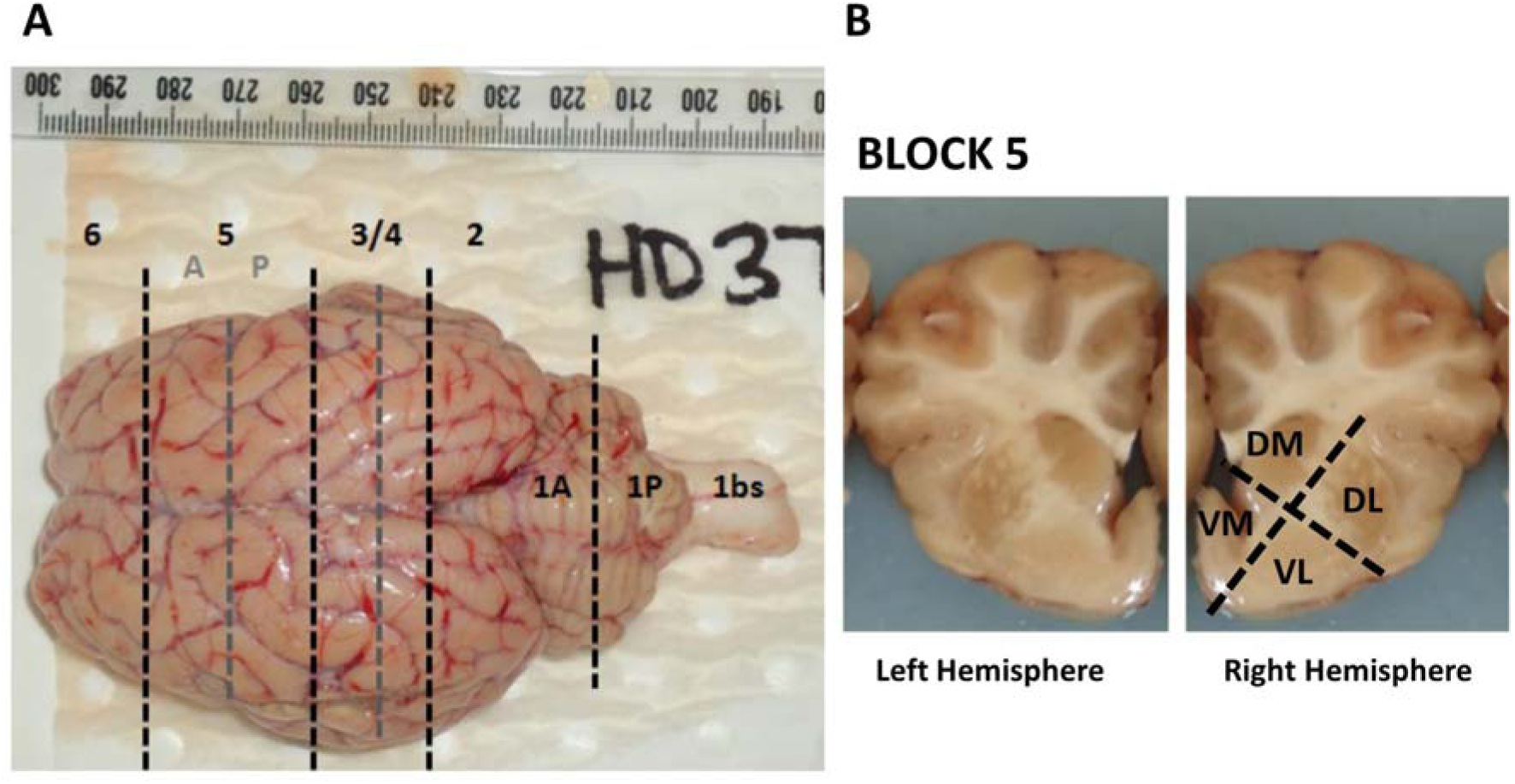
Overview of brain sectioning procedure during sample collection. (A) Superior view of a freshly extracted sheep brain showing positions of coronal sectioning. Black dashed lines indicate the initial blocks made from the whole brain. Grey dashed lines indicate sections made from the initial blocks. Block 5 was further subdivided into 5A (anterior) and 5P (posterior). (B) Left and right hemisphere of block 5 sectioning. The striatum was identified within block 5 and further subdivided into 4 subsamples, denoted as dorsal-lateral (DL), dorsal-medial (DM), ventral-medial (VM) and ventral-lateral (VL). The ventral medial sample from the anterior striatum (5A) of the left hemisphere was utilised for the generation of single nuclei transcriptomes.

### Immunohistochemical densitometric analysis

Immunohistochemical staining was performed on free floating tissue sections from the striatum of this cohort (6 *OVT73* and 6 control animals). Sections were prepared as a rostral section containing the striatum and a caudal section containing the separate caudate and putamen (Figure 2). Free floating sections were initially washed for 20 minutes in a solution of 50% methanol and 1% H_2_O_2_ to expose binding sites and block endogenous peroxidase activity. Sections were incubated in 1% NGS immunobuffer and the corresponding reagent as follows, primary GABA_A_α1 antibody (Millipore) [31] for 48 hours at 4°C; goat anti-mouse IgG secondary antibody (ThermoFisher Scientific) for 24 hours at room temperature and avidin-biotin-horseradish peroxidase (ThermoFisher Scientific) for 4 hours at room temperature. Sections were then incubated in 0.05% diaminobenzidine tetrahydrochloride (DAB) in 0.4M phosphate buffer with the addition of 40 μL/mL of 1% nickel ammonium sulphate and 0.01% H_2_O_2_. Sections were incubated until a colour change was observed. Sections were mounted onto slides and visualised on the following day.

**Figure 2.**
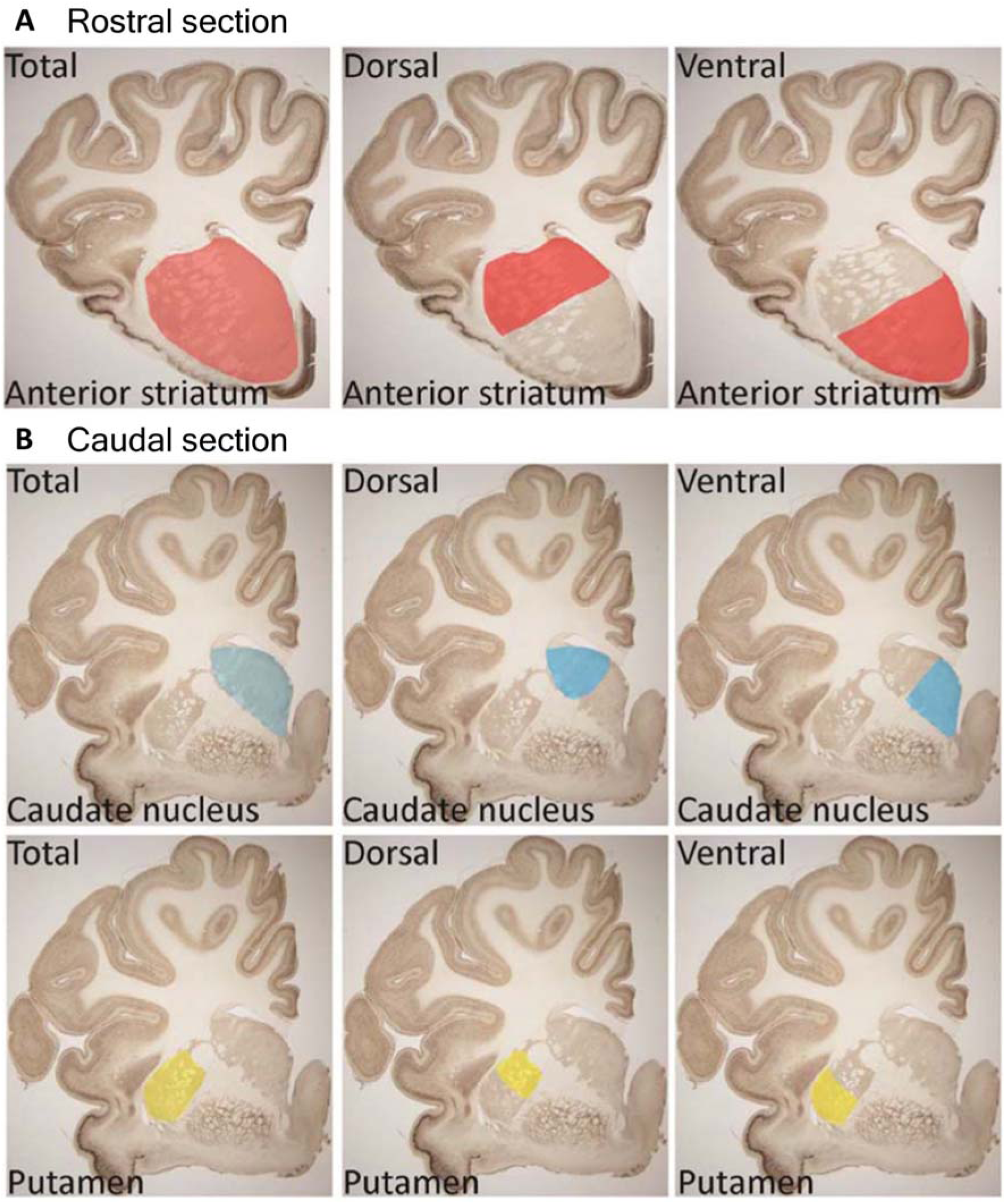
Regions of the striatum considered for immunohistochemical densitometric analysis. Ovine brain sections were prepared as a rostral section containing the striatum (A) and a caudal section containing the caudate and putamen (B).

Densitometric analysis of GABAα1 immunoreactivity was performed using a Zeiss Vslide scanner microscope (Metasystems). Images were acquired using a 20x objective. Approximately 1000 images were collected and considered for densitometric image analysis using the MetaMorph^TM^ software (Molecular Devices). Analysis was performed for the dorsal, ventral and total striatum of the rostral section and the dorsal, ventral and total caudate nucleus and putamen of the caudal section (Figure 2). A one-way analysis of variance (ANOVA) test between the means was performed to characterise significant differences in staining intensity between *OVT73* and control groups.

### Nuclei isolation for single nuclei RNA-seq

Nuclei were extracted from frozen anterior ventral-medial striatal (Figure 1) tissue utilising an adapted protocol from Krishnaswami et al., 2016 [32]. Briefly, approximately 50-100 mg of brain tissue was transferred to a dounce homogenizer containing 1mL homogenization buffer - HB (250 mM sucrose, 25 mM KCl, 5 mM MgCl_2_, 10 mM Tris buffer pH 8.0, 1 µM DTT, 1X protease inhibitor (Sigma), 0.4 U/*µ*L RNaseIn (ThermoFisher Scientific), 0.2 U/*µ*L SuperaseIn (ThermoFisher Scientific), 0.1% Triton X-100). Tissue was homogenised using 5 strokes with the loose dounce pestle A, followed by 10-15 strokes of tight dounce pestle B. The homogenate was filtered through a 40 *µ*m strainer into 5 mL Eppendorf tubes and centrifuged at 1000 rcf (4 °C) for 8 minutes. The supernatant was removed, and the pellet was resuspended in 250 *µ*L of HB. A 50% - 29% iodixanol gradient (OptiPrep™ Density Gradient Medium, Sigma) was prepared to allow removal of the myelin layer. 250 *µ*L of 50% iodixanol was added to the nuclei-HB mixture and slowly layered on top of 500 *µ*L of 29 % iodixanol in a new Eppendorf tube. The resultant gradient was centrifuged at 13,000 rcf (4°C) for 40 minutes. The supernatant and myelin were removed, and the purified nuclei pellet was resuspended in a solution containing 1 mL PBS, 1% BSA and 0.2 U/*µ*l RNAse inhibitor. 5 *µ*L of resuspended nuclei was stained with 5 *µ*L trypan blue and the quality and number of nuclei was assessed using the Countess II FL Automated Cell Counter (ThermoFisher Scientific). To reduce the cost per library, nuclei suspensions from different samples were pooled at equal concentrations in groups of 2 (Supplementary Figure 1) prior to library preparation and demultiplexed as described below.

### Library preparation and single nuclei RNA sequencing

The droplet-based Chromium methodology from 10X Genomics was utilised for the generation of single nuclei libraries. Libraries were prepared according to the Chromium Next GEM Single Cell 3’ Reagent Kits v3.1 as per manufacturer’s instructions. The single nuclei RNA-seq libraries were sequenced on the HiSeq XTen platform. Alignment of reads was performed using the CellRanger v7.0.0 pipeline with STAR v2.7.2a to the sheep Oar_rambouillet_v1.0 reference genome and annotation (Ensembl release 107). Summary statistics for single nuclei RNA-seq libraries are shown in Supplementary Figure 2.

### Sample demultiplexing

Demultiplexing of pooled nuclei associated barcodes in the CellRanger computed alignments utilized the genetic variation between individual samples. In brief, regions of the transcriptome with high read coverage (>50) were identified using the featureCounts function from the Subread package [33, 34]. These regions were used as input into Freebayes variant caller to find genomic variants for each barcode. A filtering step was subsequently applied using BCFtools [35] to remove low confidence variant calls (QUAL score < 30). Barcodes were assigned to sample ID by genotype at variant loci with the scSplit algorithm [36]. scSplit employs a hidden state model to assign nuclei associated barcodes from the pooled sample to respective groups based on an expectation-maximisation framework [36]. scSplit input parameters included an expected number of mixed samples of 2 and an estimated doublet percentage of 4% (scSplit -n 2 -d 0.04). scSplit demultiplexing outcome of barcodes is available in Supplementary File 1. The filtered unique molecular identifiers (UMI) feature barcode matrices generated by CellRanger were split according to the demultiplexed barcodes.

### Quality control and cell clustering

The filtered unique molecular identifiers (UMI) feature-barcode matrices were processed with ICARUS software [37, 38] developed by our group which utilizes the Seurat v4.0 R package [39]. A quality control filter was applied to remove low quality nuclei with gene counts less than 200 or more than 7,500. Additionally, nuclei with mitochondrial reads (>5%) were removed. From the 12 samples, a total of 28,234 high quality nuclei were recovered with an average of 1,357 median genes per nucleus (range: 1,292 – 2,607), an average of 2,672 median unique molecular identifiers (UMIs) per nucleus (range: 1,569 – 3,877) and an average percentage of transcripts originating from mitochondrial genes of 1.04% (range: 0.5% - 2.02%) (Table 1 & Supplementary Figure 3).

**Table 1.**
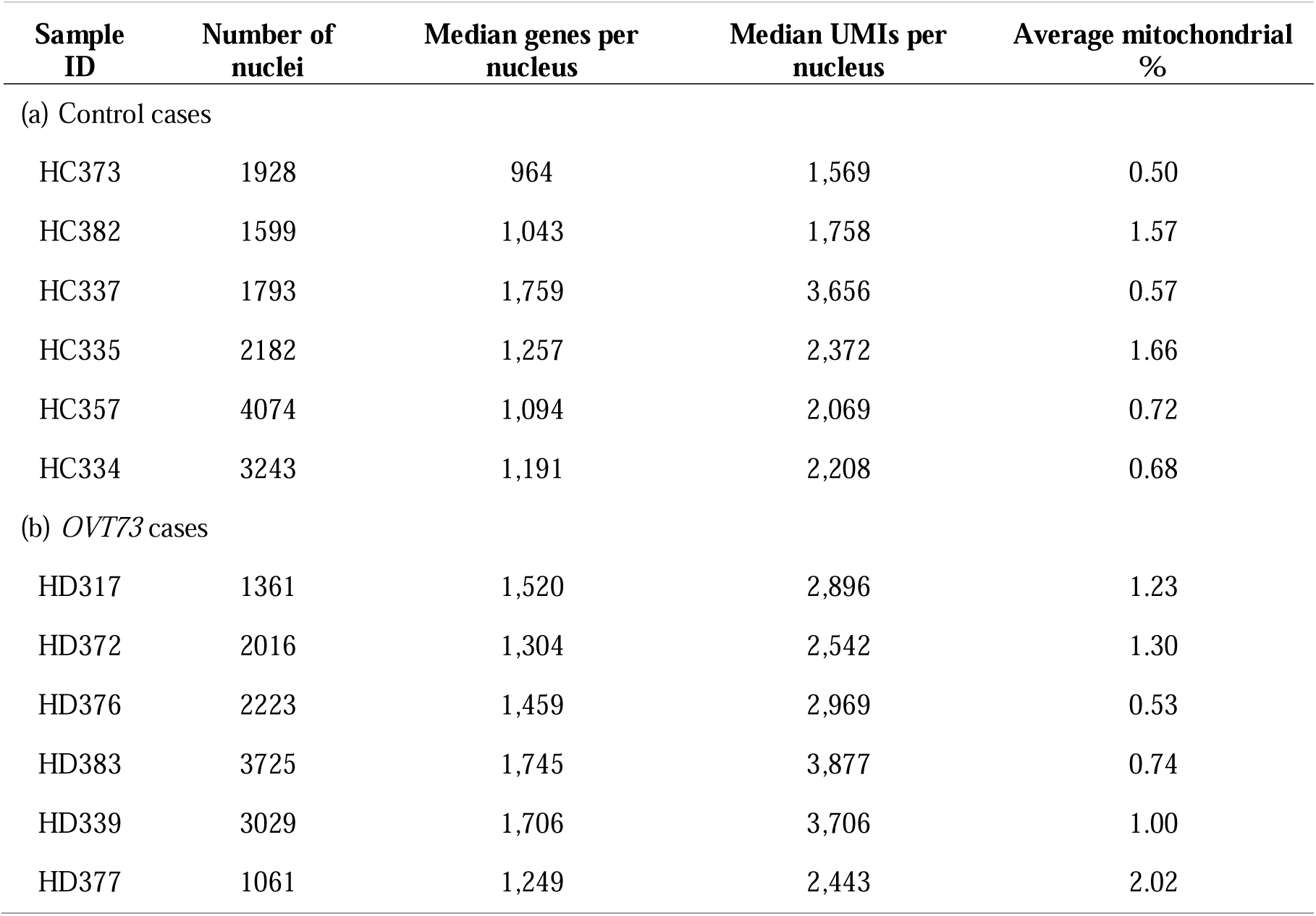
Post-quality control overview of single nuclei RNA-seq of the *OVT73* striatum.

| Sample ID | Number of nuclei | Median genes per nucleus | Median UMIs per nucleus | Average mitochondrial % |
| --- | --- | --- | --- | --- |
| (a) Control cases |  |  |  |  |
| HC373 | 1928 | 964 | 1,569 | 0.50 |
| HC382 | 1599 | 1,043 | 1,758 | 1.57 |
| HC337 | 1793 | 1,759 | 3,656 | 0.57 |
| HC335 | 2182 | 1,257 | 2,372 | 1.66 |
| HC357 | 4074 | 1,094 | 2,069 | 0.72 |
| HC334 | 3243 | 1,191 | 2,208 | 0.68 |
| (b) *OVT73* cases |  |  |  |  |
| HD317 | 1361 | 1,520 | 2,896 | 1.23 |
| HD372 | 2016 | 1,304 | 2,542 | 1.30 |
| HD376 | 2223 | 1,459 | 2,969 | 0.53 |
| HD383 | 3725 | 1,745 | 3,877 | 0.74 |
| HD339 | 3029 | 1,706 | 3,706 | 1.00 |
| HD377 | 1061 | 1,249 | 2,443 | 2.02 |

The read counts were normalised and scaled using the NormalizeData function in Seurat with parameters normalisation.method = LogNormalize and scale.factor = 10,000. Dimensionality reduction was performed on normalised read counts using a set of 3,000 top variable genes identified through the FindVariableFeatures function in Seurat. All sheep datasets (6 *OVT73* and 6 Control) were integrated using harmony [40] and cell clustering was performed with the first 30 dimensions, k-nearest neighbour value of 20 and the Louvain community detection algorithm. Cell type annotation was performed by comparison of cluster marker genes identified using the FindAllMarkers function in Seurat with parameters min.pct = 0.25 and logfc.threshold = 0.25 (Supplementary File 2 for list of cell type marker genes) against known striatal cell type markers identified in published single cell RNA-seq datasets [41–43]. Medium spiny neurons were classified into 3 sub-categories (D1, D2 and Eccentric) using cell markers described in the DropViz atlas [42].

### Differential expression analyses and Gene Ontology enrichment

Differential expression analyses were conducted for each cell type separately between *OVT73* cases and controls and also as an aggregate of all cell types (all *OVT73* nuclei compared against all control nuclei). A Wilcoxon rank sum test was performed to identify differentially expressed genes (DEGs). Genes with a p-value adjusted for false discovery rate (FDR < 0.05) and expressed in at least 10% of nuclei from either condition (*OVT73* or control) were considered to be differentially expressed. A Gene Ontology (GO) gene set enrichment analysis was performed on differentially expressed genes using the gseGO function in the R package ClusterProfiler [44], which identifies enriched GO terms using Fisher’s exact test. A ranked list of DEGs ordered by log2 fold change was used as input into gseGO. GO terms were extracted from the NCBI annotation of *Ovis aries*, retrieved with record AH107722 with the R package AnnotationHub [45]. For GO terms, a normalised enrichment score [44] was computed which reflects the overrepresentation of gene sets at the top or bottom of the gene list ordered by log2 fold change. A positive enrichment score indicates an overall upregulation of genes in the GO terms, while a negative enrichment score indicates overall downregulation of genes in the GO terms. An FDR adjusted p-value of 0.05 was set as the cut-off criteria for enriched GO terms.

### Gene co-expression analysis

Co-expression analysis was performed for the 3,000 most variable genes (identified by the FindVariableFeatures function in Seurat) across all *OVT73* and control samples. MEGENA (v1.5) was used to extract significant gene interactions and construct co-expression gene modules [46]. Default parameters detailed in the MEGENA vignette were used, including the use of Pearson’s correlations, FDR threshold of 0.05, module and hub significance p-values of 0.05 and a minimum module size of 10 genes. Modules were visualised graphically using Cystoscape v3.9.1 [47]. Module activity in cell types were determined by computing the module eigengene (first principal component) using normalised expression values of module genes. Higher eigengene values indicate higher gene expression of module genes within the cell type. The WGCNA package was used to compute module eigengenes with the moduleEigengenes function. Differential module activity between *OVT73* and control cell types was computed as a subtraction of module eigengenes of the two conditions. A randomised permutation test was used with 2,000 permutations to identify significant differential module activity between *OVT73* and control cell types.

Additionally, Gene Ontology analysis of module genes was performed using the moduleGO function in the DGCA R package with a p-value threshold of 0.05. To enhance biological interpretation, redundancy of GO terms were reduced by creating a semantic similarity matrix using the mgoSim function from the GOSemSim R package [48] and the redundancy reduced based on their semantic similarity using the reduceSimMatrix function in the R package RRVGO. Gene sets in each module were examined for its enrichment of *OVT73* versus control differentially expressed genes. Gene set enrichment analysis was performed using the GSEA function in the clusterProfiler v4.6.0 R package [44]. A ranked list of DEGs by log2 fold change was used as input into GSEA. A minimum and maximum gene set size of 10 and 500 was used with adjusted p-value FDR threshold of 0.05.

### Gene regulatory network analysis

Examination of gene regulatory networks for cell types across *OVT73* and control samples was conducted using R SCENIC v1.2.0 [49]. SCENIC performs cis-regulatory transcription factor binding motif analysis on a set of co-expressed transcription factors and genes. Transcription factor genes were defined from the curated list by Lambert et al., [50]. Given, a public sheep transcription factor motif database was not available, a custom transcription factor motif database for sheep was made for this study. The motif database was generated as follows, the 500bp sequence upstream and 100bp sequence downstream of the transcriptional start site (TSS) for each gene was extracted from the oar_rambouillet v1.0 reference genome (ensemble version 107). Additionally, the 10kb sequence upstream and downstream of the TSS were also extracted. The files are available on zenodo (https://doi.org/10.5281/zenodo.8057929) under oar_rambouillet_v1.0_500bp_up_100bp_down.fa and oar_rambouillet_v1.0_10kbp_up_10kbp_down.fa. These regions were assessed for transcription factor binding motifs based on the 2022 SCENIC+ motif collection (https://resources.aertslab.org/cistarget/motif2tf/) using the create_cistarget_motif_databases.py function of the create_cisTarget_databases package (https://github.com/aertslab/create_cisTarget_databases). Code used to generate database is shared under https://doi.org/10.5281/zenodo.8057929.

A normalised gene expression matrix of DEGs identified between *OVT73* and controls for each cell type separately and also in the aggregate analysis (all *OVT73* nuclei compared against all control nuclei) were used as inputs into SCENIC. The full lists of DEGs are available in Supplementary File 3. Co-expression modules were constructed using GENIE3 [49, 51] and transcription factor motifs were scored using RcisTarget v1.18.2. Transcription factor-regulated gene modules (regulons) with 10 or more genes were considered. The activity of each regulon in *OVT73* and control cell types (regulon activity) were scored using the AUCell algorithm which computes the enrichment of regulons as an area under the recovery curve across a ranking of all genes in a cell based on their normalised expression values. A high regulon activity indicates genes within the regulon are positively regulated by the transcription factor. Regulons were visualised graphically using Cystoscape v3.9.1 [47]. Differential regulon activity between *OVT73* and control cell types was computed as a subtraction of regulon activity of the two conditions. A randomised permutation test was used with 2,000 permutations to identify significant differential regulon activity between *OVT73* and control cell types. Additionally, a regulon specificity score (RSS) was determined which employs the Jensen-Shannon divergence metric to assess cell-type specificity of regulon activity [49]. Regulons in cell types with a specificity score of 1 indicates exclusive expression of the regulon in that one cell type, while a specificity score of 0 indicates the regulon is evenly expressed across all cell types.

### Cell-cell communication analysis

The inference of cell signalling crosstalk through the transcription levels of ligand receptor pairs was examined using CellChat. CellChat incorporates a comprehensive database of ligand receptor interactions, soluble agonists/antagonists and stimulatory/inhibitory membrane bound co-receptors to infer cell-cell communications from single cell RNA-seq data based on mass action models, social network analysis tools and pattern recognition methods [52]. In brief, intercellular communications were inferred through 3 steps; (1) identification of differentially expressed ligands and receptors genes for each cell type using the Wilcoxon rank sum test with a p-value threshold of 0.05. (2) A communication probability (interaction strength) is computed by modelling ligand-receptor interactions using the law of mass action on average expression values of a ligand in one cell group and a receptor of another cell group. This calculation accounts for the number of cells within each cell group. (3) Significant interactions are identified using a permutation test that randomly permutes cell group labels and recomputes the interaction probability.

A comparison of cell-cell communication in *OVT73* versus control cell types was performed using default parameters detailed in the CellChat v1.6.1 vignettes. Ligand-receptor interactions were extracted from the human CellChat database. Interactions involving 50 or less nuclei were filtered out using the filterCommunication function. Quantification of similarity among intercellular communication networks was performed using the functional similarity measure as described by Jin et al., 2021 [52] and plotted onto a two-dimensional manifold.

## Results

### Upregulation of genes involved in synaptic transmission in *OVT73* medium spiny neurons

Following quality control, a total of 28,234 nuclei were recovered from the striatum of 6 *OVT73* sheep and 6 age matched wild-type controls (Table 1). A total of 13 cell types were identified including 12,507 oligodendrocytes (expressing marker genes *MOG* and *PLP1*), 6,093 medium spiny neurons (*RGS9*, *PDE10A*), 3,024 oligodendrocyte precursor cells (*PCDH15*, *PDGFRA*), 2,601 microglia (*CSF1R*, *CX3CR1*), 2,181 astrocytes (*AQP4*, *GFAP*), 1,068 neuroblasts (*DCX*, *ADARB2*), 683 interneurons (*ELAVL2*, *CLSTN2*) and 77 endothelial cells (*FLT*, *MECOM*). Interneurons were subcategorised based on markers defined in Munoz Manchado et al., [53] into 526 PV/Th interneurons (*HS3ST2*, *PTHLH*), 107 SST/NPY interneurons (*SST*, *NPY*) and 50 cholinergic interneurons (*CHAT*). Medium spiny neurons (MSN) were subcategorised according to cell markers defined in Saunders et al., [42] into 2,776 D1 medium spiny neurons (*TAC1*, *EBF1*), 2,998 D2 medium spiny neurons (*DRD2*, *PENK*) and 319 eccentric medium spiny neurons (*OTOF*, *FOXP2*). Full cell marker lists are provided in Supplementary File 2, and cell type distributions shown in Figure 3C. Across all cell types, we observed a significant reduction in the proportion of oligodendrocytes in the *OVT73* derived tissue compared with control cases (ANOVA, p = 1.24×10^-5^).

**Figure 3.**
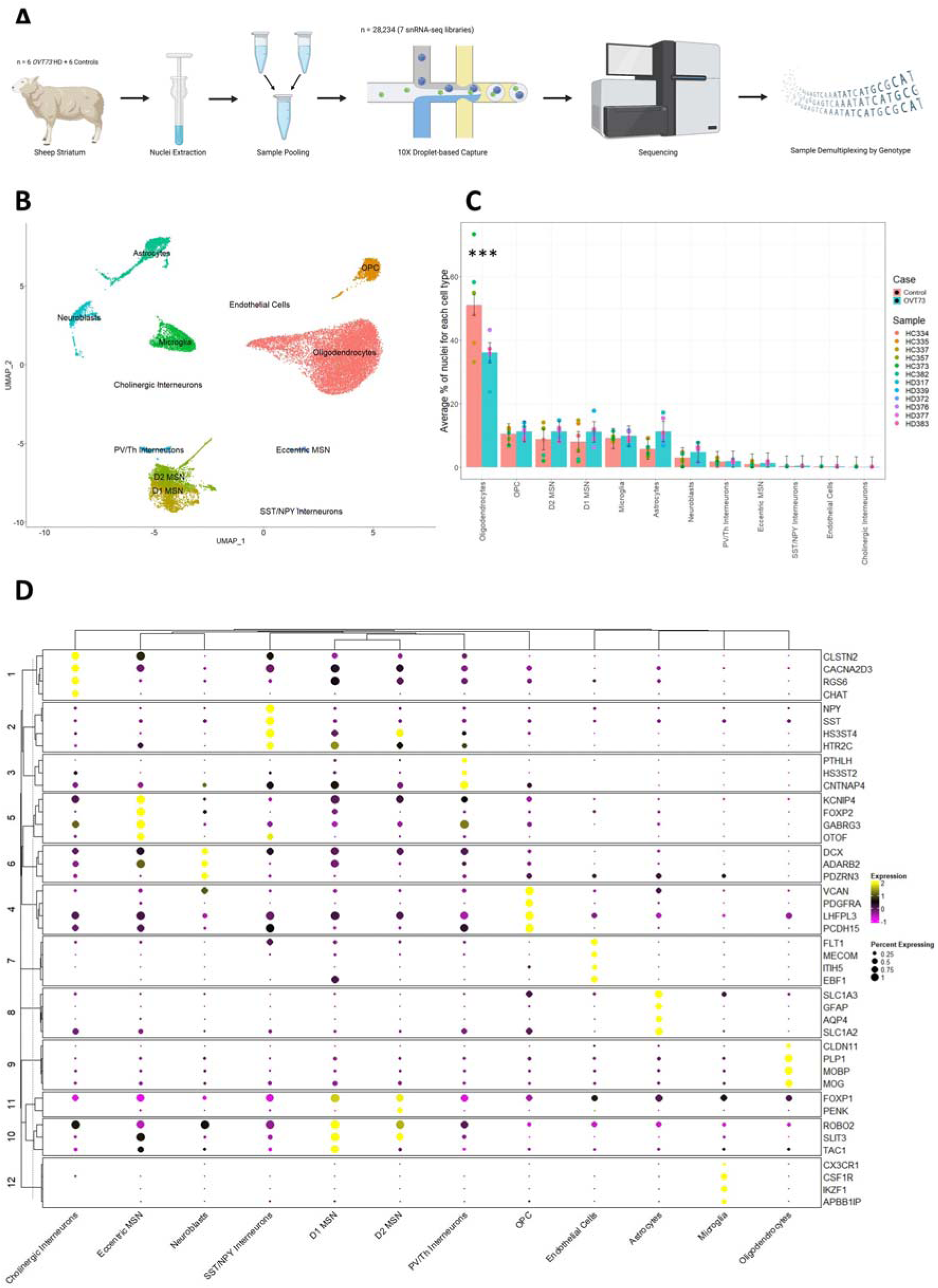
Single nuclei RNA-seq of the *OVT73* sheep striatum. (A) Experimental design. Nuclei were extracted from the striatum of 12 sheep (6 *OVT73* and 6 controls) and pooled to form 7 nuclei suspensions. Single nuclei RNA libraries were generated from the sample multiplexed suspensions and sequenced. Reads were demultiplexed based on the natural genetic variation between pooled samples. Panel image created with BioRender.com. (B) UMAP visualisation of clusters identified and annotated by cell marker gene expression. (C) Proportions of identified cell types across the 12 animals. A significant decrease in oligodendrocytes between *OVT73* and control cases was observed (***ANOVA, p < 0.001) (D) Clustered dot plot of selected cell-type enriched markers for annotated cell types in the sheep striatum.

To investigate differential gene regulation in the *OVT73* HD striatum, differential expression analyses were conducted between *OVT73* and controls for each cell type separately and also as an aggregate of all cell types (all *OVT73* nuclei compared against all control nuclei). A gene was considered to be differentially expressed if it presented with a p-value adjusted for false discovery rate less than 0.05 that was also expressed in at least 10% of nuclei from either *OVT73* or control. The ratio of differentially expressed genes (DEGs) identified to the median number of expressed genes per nucleus was 5.08 (6,748 DEGs/1,329 median number of expressed genes of all nuclei) for the aggregate analysis. When examining DEGs in individual cell types, we observed a ratio of 2.14 in oligodendrocytes (2,652 DEGs/1,238 median number of expressed genes of oligodendrocyte nuclei), 0.71 in OPCs (1,286/1,812), 1.57 in D2 medium spiny neurons (7,095/4,516), 0.69 in D1 medium spiny neurons (3,329/4,850), 0.10 in eccentric medium spiny neurons (177/1,687), 0.80 in microglia (837/1,049), 0.71 in neuroblasts (632/895), 0.71 in astrocytes (1,191/1,687) and 0.07 in PV/Th interneurons (310/4,398) (Figure 4B). No DEGs were detected in endothelial cells, cholinergic interneurons, and SST/NPY interneurons, likely due to their small population sizes. The statistical power to detect differences in gene expression is dependent on the number of cells, consistent with the highest proportion of DEGs being observed in oligodendrocytes, with the highest number of cells. When comparing D1 medium spiny neurons, D2 medium spiny neurons, OPC, microglia, astrocytes and neuroblasts with similar cell numbers, a greater proportion of DEGs was observed for D2 medium spiny neurons indicative of an increased transcriptomic variability for this cell type in the *OVT73* animals. The full list of DEGs is available in Supplementary File 3 and volcano plots for individual cell type DEG analysis are available in Figure 4A and Supplementary Figure 4.

**Figure 4.**
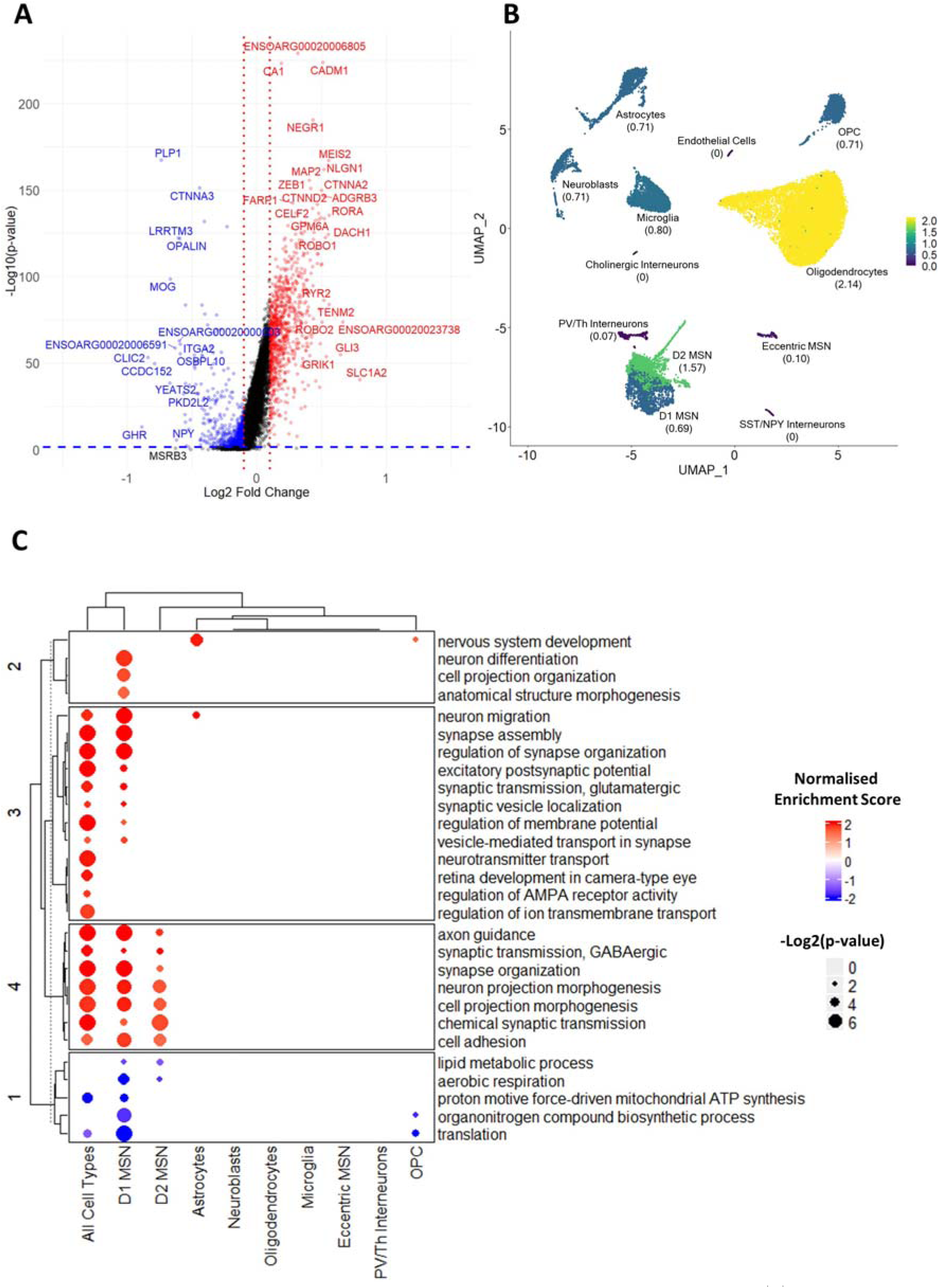
Differentially expressed genes in *OVT73* versus control sheep striatum. (A) Volcano plot of differentially expressed genes (DEGs) identified from the aggregate analysis (all *OVT73* nuclei vs all control nuclei regardless of cell type). Horizontal blue line shown at p=0.05, vertical red lines shown at log2 fold change of -0.1 and 0.1. (B) Proportion of DEGs over the median number of expressed genes in each cell type. (C) Dot plot of most significant Gene Ontology terms (ordered by lowest FDR adjusted p-values) in cell types. Dots were coloured by normalised enrichment score. Normalised enrichment scores reflect the overrepresentation of gene sets at the top or bottom of a gene list ranked by log2 fold change. A positive enrichment score indicates upregulation of genes in the GO terms, while a negative enrichment score indicates downregulation of genes in the GO terms. The size of the dot represents the FDR adjusted p-value. A false discovery rate (FDR) adjusted p-value of 0.05 was set as the cut-off criteria for enriched GO terms.

The top enriched Gene Ontology (GO) terms (ordered by FDR adjusted p-values) associated with DEGs in the aggregate analysis included synapse assembly and organisation, GABAergic and glutamatergic synaptic transmission, axon guidance, neuronal projection morphogenesis, regulation of ion transport, aerobic respiration, and proton motive force driven mitochondrial ATP synthesis (Figure 4C). Further refinement of these results revealed differential expression in D1 and D2 medium spiny neurons were the most significant contributors to the enrichment of the above GO terms. D1 and D2 medium spiny neurons are the early affected cell type in HD. Interestingly there was an overall upregulation of the genes underlying the GO terms, synapse assembly and organisation, GABAergic and glutamatergic synaptic transmission, axon guidance, neuronal projection morphogenesis and regulation of ion transport. There was also a down regulation of transcription from genes underlying the GO terms, aerobic respiration and proton motive force-driven mitochondrial ATP synthesis. The full list of enriched GO terms is available in Supplementary File 4.

### Co-expressed gene modules involved in synaptic transmission show increased activity in *OVT73* medium spiny neurons and astrocytes

To identify and investigate sets of genes that were co-expressed in the *OVT73* versus control cell types, we performed co-expression module analysis with MEGENA. We identified several gene modules involved in synaptic transmission that exhibited higher module activity in *OVT73* medium spiny neurons and *OVT73* astrocytes compared to controls. Module activity was determined by computing the module eigengene (first principal component) using normalised expression values of module genes. A module centred around *DLGAP2*, *GRIN2B*, *CACNA1C*, *CACNA1B*, *CNTNAP5*, *SYT1*, *KCTD16*, and *GRIN2A* genes (Module 2, M2) showed higher module activity in *OVT73* D1 medium spiny neurons compared to control D1 medium spiny neurons (differential module activity of 0.054; p < 0.0005, randomised permutation test with 2,000 permutations) and *OVT73* D2 medium spiny neurons compared to control D2 medium spiny neurons (differential module activity of 0.06, p < 0.0005) (Figure 5B, Supplementary Figure 5 and Supplementary Figure 6). The most significant GO terms for module genes included synapse organisation, synapse assembly, chemical synaptic transmission, dicarboxylic acid catabolic process, glutamate metabolic process and regulation of synaptic vesicle exocytosis (Figure 6). Additionally, this module was enriched for D1 medium spiny neurons DEGs (gene set enrichment analysis, normalised enrichment score = 0.32, FDR = 0.0077) and D2 medium spiny neurons DEGs (normalised enrichment score = 0.41, FDR = 7.75×10^-5^) which included DEGs encoding glutamate receptors complexes (*GRIN2A*, *GRIN2B*, *FRRS1L*, *CNIH3, GRIK5*), vesicle transport proteins (*SNAP25*, *SYT1*, *SPTBN4, SYNPR*, *SYN1*) and post-synaptic density proteins (*CLSTN2*, *DLGAP2, DLGAP4*, *RIMS3*).

**Figure 5.**
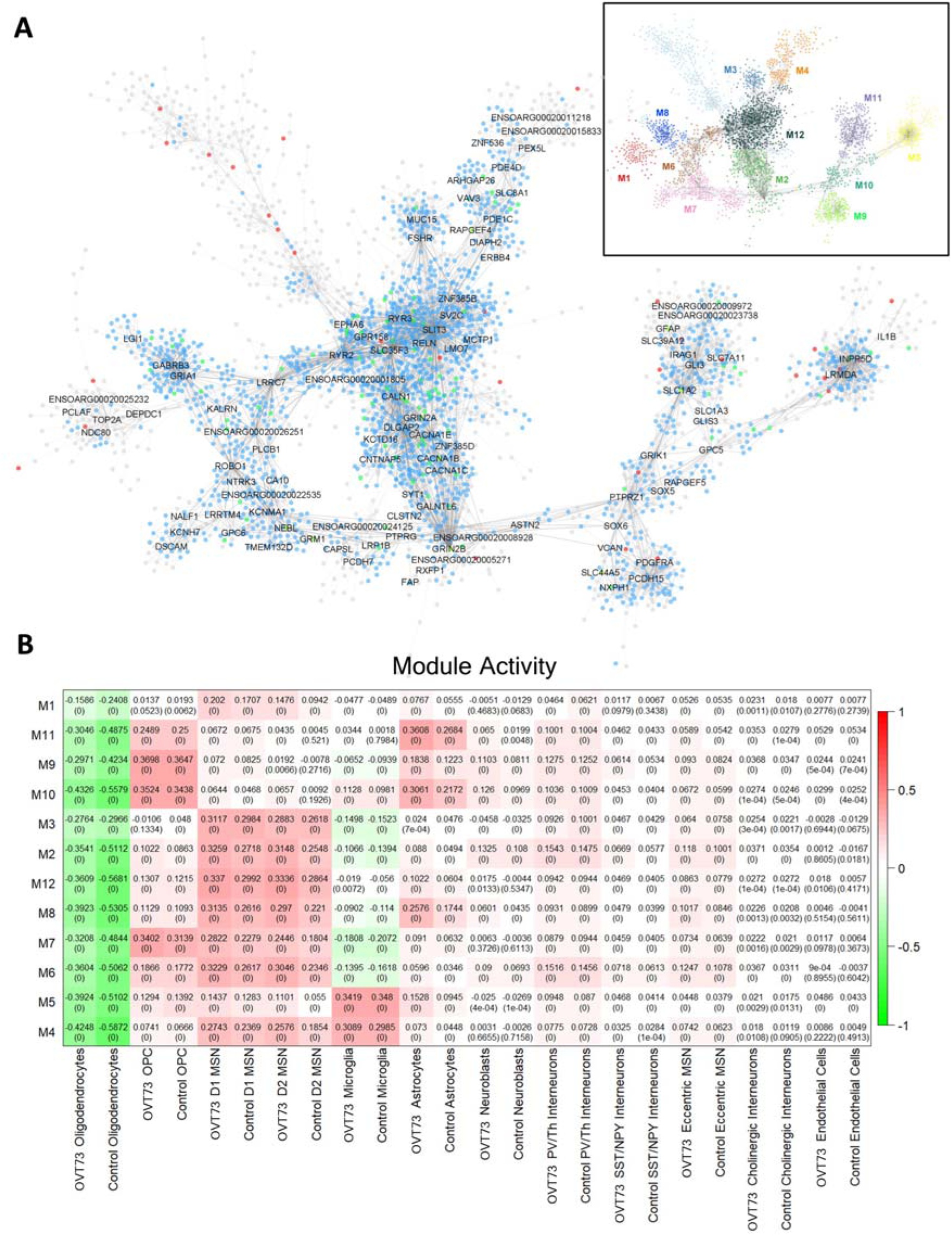
Co-expression gene modules. (A) Co-expression modules generated using the Multiscale Embedded Gene Co-Expression Network Analysis (MEGENA). A total of 12 modules were identified with the structure outlined in the top right. Module hub genes are labelled with connected genes represented as dots. Genes differentially expressed between *OVT73* and controls are coloured in blue. Genes with evidence to support an interaction with *HTT* as curated by the HDinHD database [54] were coloured in red. Genes that were both differentially expressed and a known *HTT* interactor were coloured in green. (B) Co-expression module activity in *OVT73* and control cell types. Module activity in cell types were determined by computing the module eigengene (first principal component) using normalised expression values of module genes. Eigengene values are shown with p-values of the correlation shown in parentheses in each square. Higher eigengene values indicate higher gene expression of module genes within the cell type.

**Figure 6.**
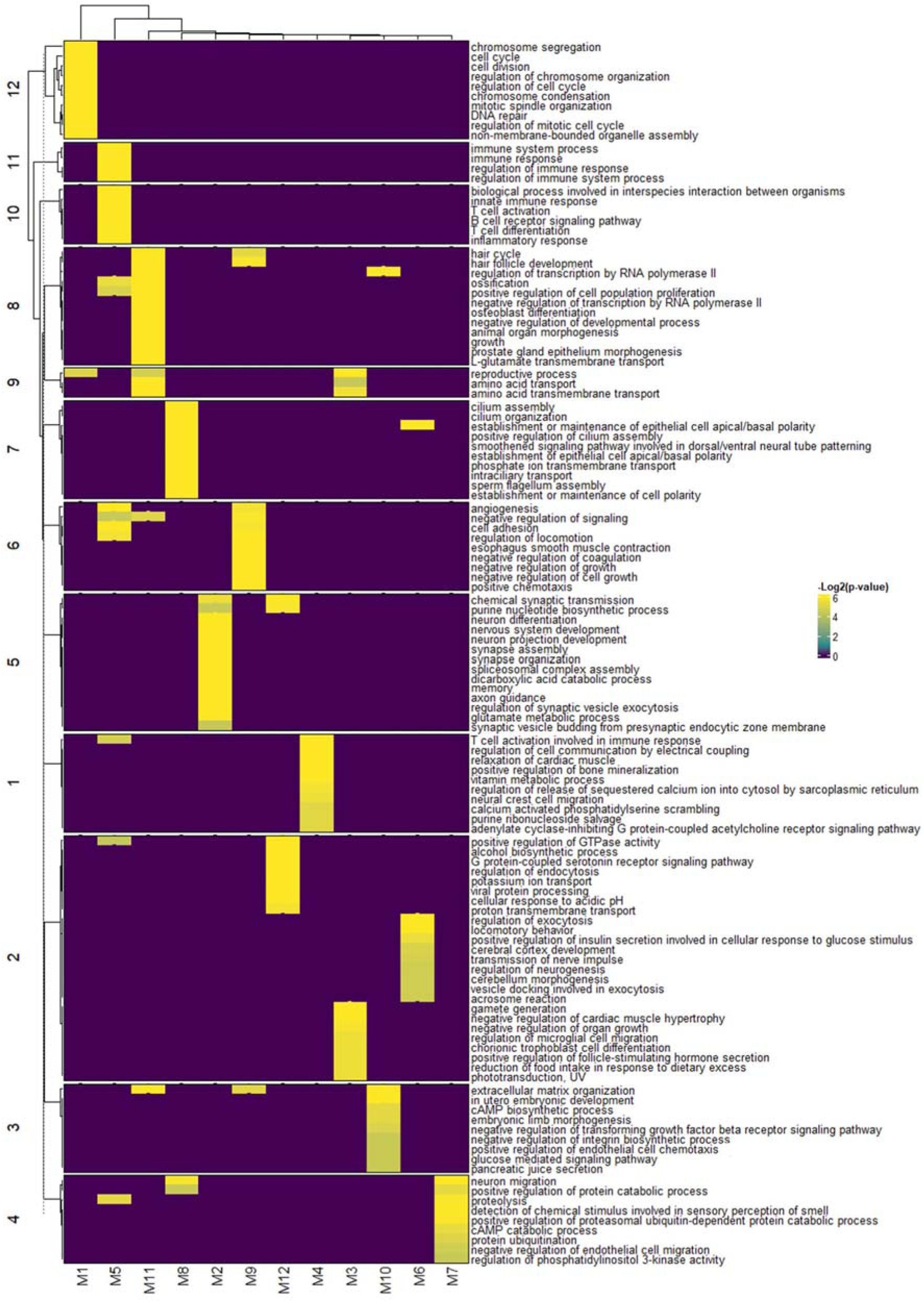
Clustered heatmap of gene ontology terms for gene modules. The 10 most significant gene ontology terms ordered by FDR adjusted p-values for each gene module are shown. Enriched GO terms with FDR < 0.05 are shown.

Another module defined by genes, *ROBO1*, *LRRC7*, *KALRN*, *PLCB1* and *NTRK3* (Module 6, M6) also showed higher module activity in *OVT73* D1 medium spiny neurons compared to control D1 medium spiny neurons (differential module activity of 0.061, p < 0.0005) and OVT73 D2 medium spiny neurons compared to control D2 medium spiny neurons (differential module activity of 0.07, p < 0.0005) (Figure 5B, Supplementary Figure 5 and Supplementary Figure 6). Most significant GO terms associated with this module included regulation of exocytosis, vesicle docking involved in exocytosis and transmission of nerve impulse (Figure 6). This module was also enriched for D1 medium spiny neurons DEGs (normalised enrichment score = 0.55, FDR = 1.32×10^-7^) and D2 medium spiny neurons DEGs (normalised enrichment score = 0.43, FDR = 0.0035) and included DEGs involved in exocytosis (*SYT10*, *KALRN*), synaptic transmission (*RIMS2*, *RIMBP2*, *STXBP1*, *DLG3*) and neuronal projection (*ROBO1*, *LRRC7*, *PLXNA4*).

A module centred around *SLC1A2*, *SLC7A11*, *GLI3*, *IRAG1*, *GFAP* and *SLC39A12* genes (Module 11, M11) was enriched in *OVT73* astrocytes compared to control astrocytes (differential module activity of 0.092, p < 0.0005). Most significant GO terms for modules genes included L-glutamate transmembrane transport and amino acid transport. The module showed an enrichment of astrocyte DEGs (normalised enrichment score = 0.57, FDR = 2.5×10^-3^) and included several genes involved in glutamate transport (*SLC1A2*, *SLC1A3*, *SLC7A11*). Full lists of genes in gene modules are available in Supplementary File 5. The full list of enriched GO terms for modules is available in Supplementary File 6.

### Cell-cell signalling is elevated in *OVT73*

The differential expression analyses and co-expression analyses both indicated that synaptic signalling and transmission may be increased in *OVT73*. This led us to examine the cell-cell communication networks using CellChat. The CellChat algorithm infers intercellular communication probabilities in single nuclei RNA-seq expression data based on differentially expressed ligand and receptor genes in each cell type [52]. Cell-cell communication analysis also supported increased synaptic signalling in neuronal and glial cells of the *OVT73* striatum. Comparison of ligand-receptor pair expression from *OVT73* and control revealed a higher total number of ligand-receptor interactions (4,030 and 3,509 detected ligand-receptor pairs for *OVT73* and control respectively) and a higher sum of all communication probabilities (200 and 130 for *OVT73* and control respectively) across all cell types in *OVT73*. The cell types that showed the greatest increase in the number of incoming (receptor to ligand) and outgoing (ligand to receptor) ligand-receptor interactions included D2 medium spiny neurons and astrocytes. Interestingly, compared to D2 medium spiny neurons, D1 medium spiny neurons exhibited a relatively lower degree of increase. The communication probability was greater in the *OVT73* dataset for all neuronal cells (D1, D2, eccentric medium spiny neurons and PV/Th, NPY/SST, cholinergic interneurons), OPCs and astrocytes indicating more cell-cell crosstalk in the *OVT73* cell state (Figure 7A & B, thicker lines indicate higher number of ligand receptor interactions or higher communication probability in *OVT73*). In contrast, no differences in cell-cell signalling were detected in microglia from *OVT73* compared to controls (Supplementary Figure 7).

**Figure 7.**
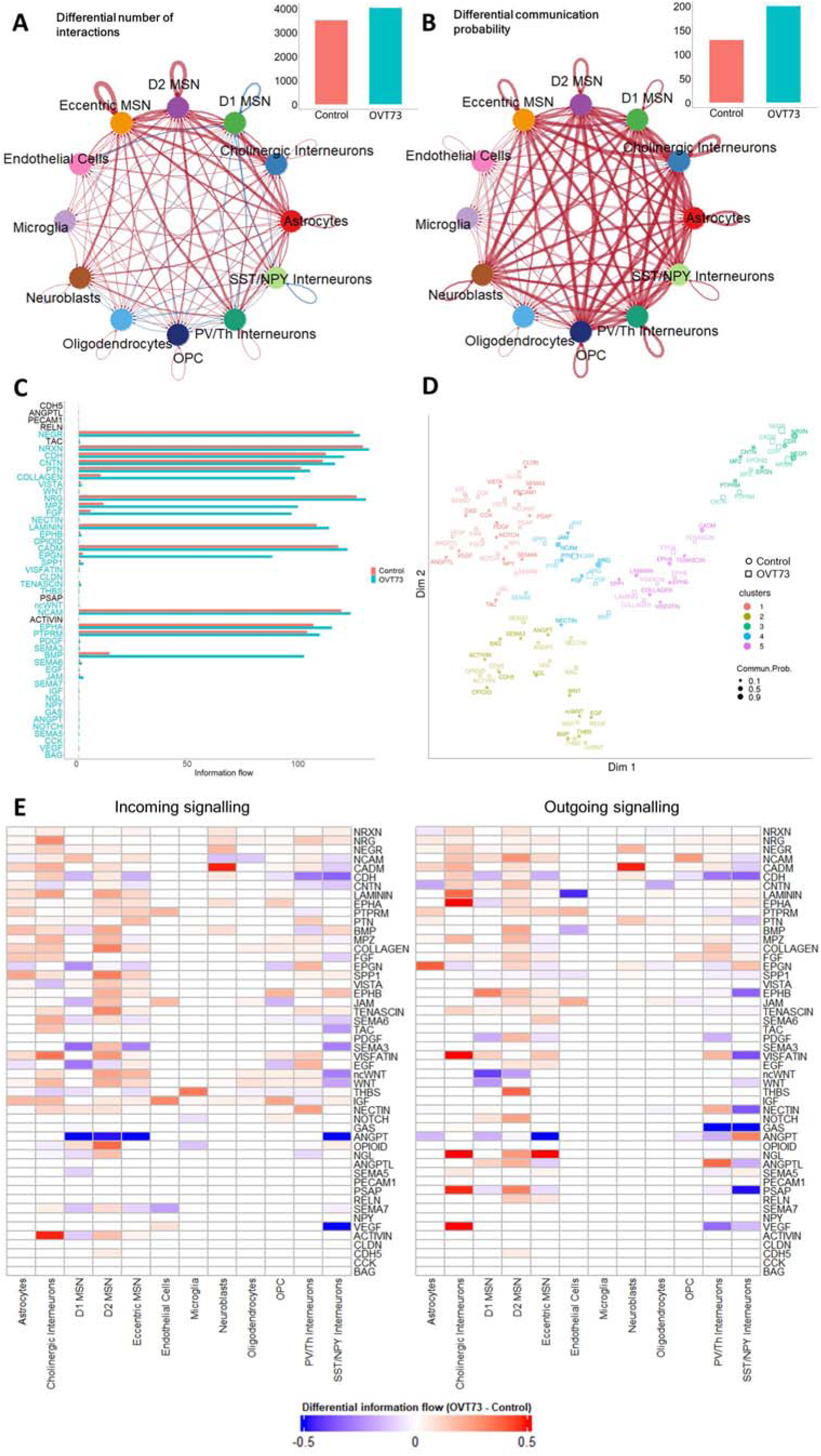
CellChat cell-cell communication networks inferred from expression of ligand-receptor pairs. Circle plot showcasing the (A) differential number of ligand-receptor interactions and (B) differential communication probability between *OVT73* and controls for any two cell types. Red arrows indicate increased number/communication probability in *OVT73*, blue arrows indicate decreased number/communication probability in *OVT73*. Thickness of line indicates greater number/communication probability. The total number of ligand-receptor interactions and total interaction strength for *OVT73* and controls is also shown as bar graphs on the top right. (C) Information flow for all signalling pathways in *OVT73* and control. The information flow for a given signalling pathway is defined as the sum of communication probabilities of all ligand receptor pairs in the pathway. (D) Two-dimensional manifold projection of signalling pathways based on its functional similarity. Signalling pathways that cluster together are likely to exhibit similar or redundant roles. Circle and square symbols represent the pathways in *OVT73* and control respectively. (E) Differential information flow between *OVT73* and control cell types for outgoing (ligand to receptor) and incoming (receptor to ligand) signalling pathways. A positive value indicates more communication in *OVT73*.

Examination of signalling pathways showed greater information flow (defined as the sum of communication probabilities of all ligand receptor pairs in the signalling pathway) in neurexin (NRXN), neuregulin (NRG), protein tyrosine phosphatase receptor M (PTPRM), contactin-1 (CNTN), neuronal growth regulator (NEGR), laminin, cell adhesion molecule (CADM), neural cell adhesion molecule (NCAM), ephrin type receptor A (EPHA) and pleiotrophin (PTN) pathways in *OVT73* D2 medium spiny neurons. Interestingly, the greatest increase in communication was observed in myelin protein zero (MPZ), collagen, bone morphogenetic protein (BMP), and fibroblast growth factor (FGF) signalling pathways (Figure 7C & 7E, Supplementary Figure 8). Clustering of the signalling pathways based on the functional similarity measure as described by Jin et al., [52] on a two-dimensional manifold revealed NEGR, NRXN, CDH, CADM, EPGN, PTPRM, MPZ, CNTN signalling pathways clustered together (cluster 3, Figure 7D), EPHA, LAMININ, COLLAGEN signalling pathways clustered together (cluster 5) and JAM, FDF, NCAM, PTN and NRG signalling pathways clustered together (cluster 4). Signalling pathways that cluster together are likely to exhibit similar or redundant roles. Individual ligand receptor pairs that showed the greatest increase in communication probability between *OVT73* medium spiny neurons and control included *NEGR1*-*NEGR1*, *NRXN1*-*NLGN1*, *CADM1*-*CADM1*, *NRXN3*-*NLGN1*, *CDH2*–*CDH2*, *NRG1*-*ERBB4* and *NRG3*-*ERBB4*. These ligand-receptor pairs have been implicated in various neurodevelopmental processes including myelination [55–58], glutamatergic and GABAergic synapse development [59, 60], neurite outgrowth and general nervous system development [61–68]. Individual ligand receptor pair interactions for astrocytes and medium spiny neurons (D1, D2) are shown in Supplementary Figure 9. Communication probabilities between all ligand receptor pairs across all cell types is available in Supplementary File 7.

### Evidence for glutamate excitotoxicity in the *OVT73* striatum

The glutamate excitotoxicity hypothesis proposes neuronal stress and eventual initiation of cell death pathways arising from elevated levels of synaptic glutamate and excessive signalling through ionotropic glutamate receptors [3–6]. We have identified evidence in support of glutamate excitotoxic stress in the *OVT73* striatum through a widespread transcriptional upregulation of ionotropic glutamate receptors (NMDA, AMPA, kainate and delta receptors) (Figure 8A). Transcription from genes encoding the NMDA receptor subunit B (*GRIN2B*), AMPA receptor subunits 1 (*GRIA1*), 2 (*GRIA2*) and 4 (*GRIA4*), kainate receptor subunits 1 (*GRIK1*), 3 (*GRIK3*), 4 (*GRIK4*) and 5 (*GRIK5*) and delta receptor 2 (*GRID2*) were upregulated in *OVT73* D2 medium spiny neurons compared to controls. *OVT73* D1 medium spiny neurons follow a similar pattern of upregulation with increased expression of genes *GRIA1*-*4*, *GRID2*, *GRIK1*, *GRIK*2 and *GRIK5*. D1 and D2 medium spiny neurons are the most vulnerable cell type in HD. We also observed an upregulation of transcription of the glutamine to glutamate conversion enzyme glutaminase (*GLS*) in *OVT73* D1 and D2 medium spiny neurons and *OVT73* astrocytes. Upregulation of *GLS* suggests increased production of glutamate which may be released into the synaptic space, bind glutamate receptors, and trigger a feedback loop to upregulate glutamate receptor expression.

**Figure 8.**
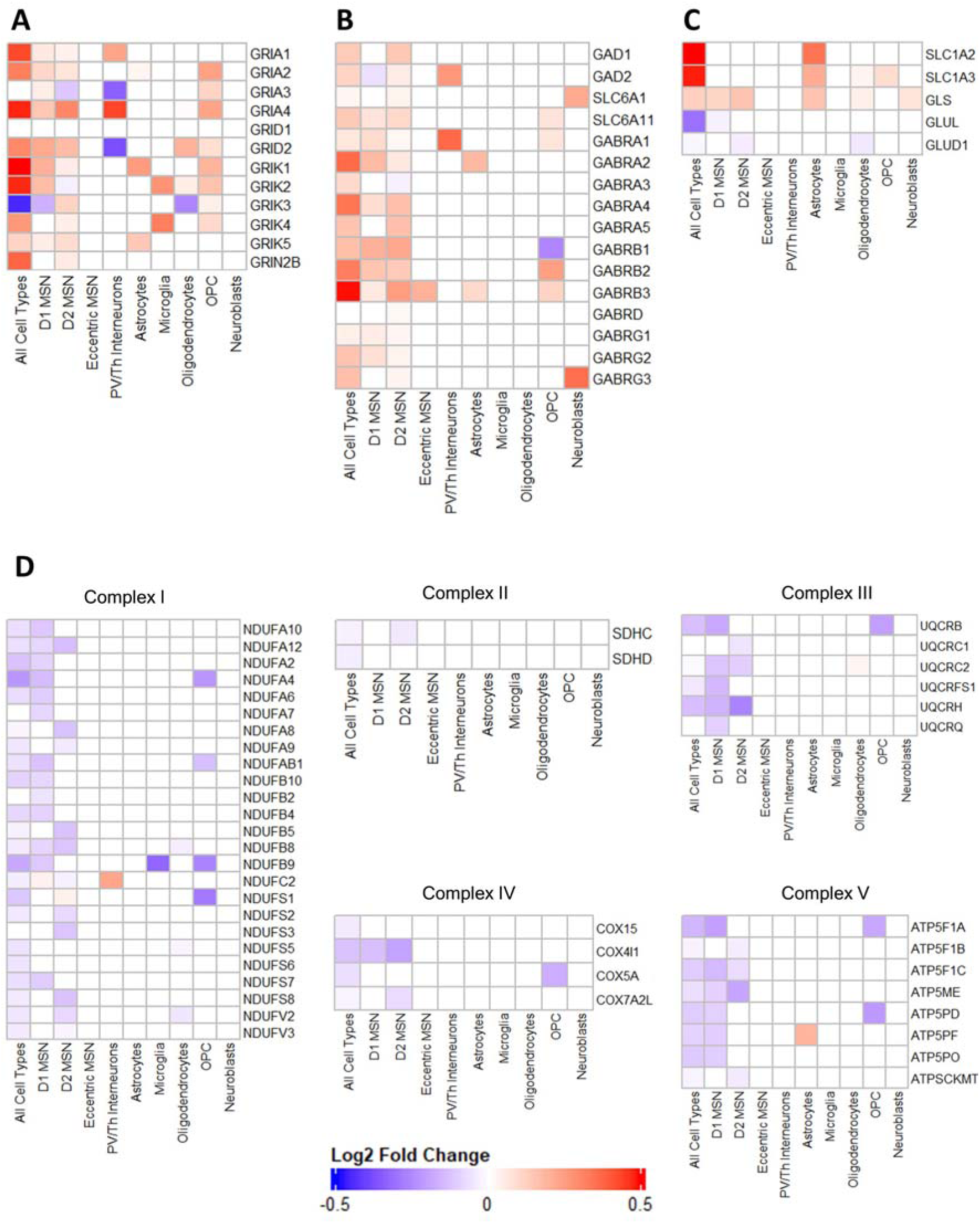
Heatmap of log2 fold change of selected differentially expressed genes between *OVT73* and control cell types implicated in glutamate excitotoxicity. Differential gene transcriptional regulation of (A) glutamate receptors (NMDAR, AMPAR, Kainate, delta receptors), (B) GABA_A_ receptors and glutamate to GABA conversion genes, (C) glutamate uptake transporters and glutamine-glutamate cycle genes and (D) genes encoding oxidative phosphorylation complexes.

Excitotoxicity is commonly observed with oxidative stress signatures represented by reduced activity of the oxidative phosphorylation complexes [69–71]. In keeping with this we have observed a downregulation of several genes encoding oxidative phosphorylation complexes including complex I (*NDUFA2,4,6,7,8,9,10,12, NDUFAB1, NDUFB2,4,5,8,9, NDUFC2, NDUFS1,3,7,8*, *NDUFV2, NDUFV3*), complex II (*SDHC*), complex III (*UQCRB, UQCRC1, UQCRC2, UQCRFS1, UQCRH, UQCRQ*), complex IV (*COX4I1, COX7A2L*) and complex V (*ATP5F1A, ATPF1B, ATP5F1C, ATP5ME, ATP5PD, ATP5PF, ATP5PO*, *ATPSCKMT*) in *OVT73* D1 and D2 medium spiny neurons. In addition, we observed transcriptional upregulation of several genes likely due to a compensatory response to glutamate excitotoxicity. The transcription of genes encoding glutamate uptake transporters GLT (*SLC1A2*) and GLAST (*SLC1A3*) were upregulated in *OVT73* astrocytes suggesting a response to remove excess synaptic glutamate (Figure 8C).

### Immunohistochemistry densitometric analysis show a region dependent reduction in GABA_A_**α**1 receptor subunit

We also describe a transcriptional upregulation of genes encoding the gamma-aminobutyric acid A receptor subunits (GABA_A_) including the α subunit (*GABRA1*, *GABRA2*, *GABRA4* and *GABRA5*), β subunit (*GABRB1*, *GABRB2*, *GABRB3*) and γ subunit (*GABRG1, GABRG2*, *GABRG3*) in *OVT73* D1 and D2 medium spiny neurons compared to controls. Additionally, an upregulation of genes encoding GABA transporters GAT1 (*SLC6A1*) and GAT4 (*SLC6A11*) and genes encoding the glutamate to GABA conversion enzymes, glutamate decarboxylase (*GAD1* and *GAD2*) was also observed in *OVT73* D2 medium spiny neurons (Figure 8B). Taken together, these observations suggested increased GABA_A_ receptor signalling is likely a result of increased glutamate conversion to GABA.

Immunohistochemical densitometric analysis of the GABA_A_ receptor subunit α1 (encoded by *GABRA1*) was performed on rostral and caudal sections of the striatum of the same *OVT73* and control animals (Figure 2). GABA_A_α1 immunoreactivity was reduced in the dorsal striatum of the rostral section (35% reduction, p-value = 0.048), dorsal caudate (35% reduction, p-value = 0.043) and dorsal putamen (39% reduction, p-value = 0.043) of the caudal section of *OVT73* animals (Figure 9, Figure 10, Supplementary Figure 10). However, no changes in GABA_A_α1 immunoreactivity were observed for ventral regions (striatum, caudate, putamen) of rostral and caudal sections which included the region of the tissue that was utilised in the single nuclei RNA-seq.

**Figure 9.**
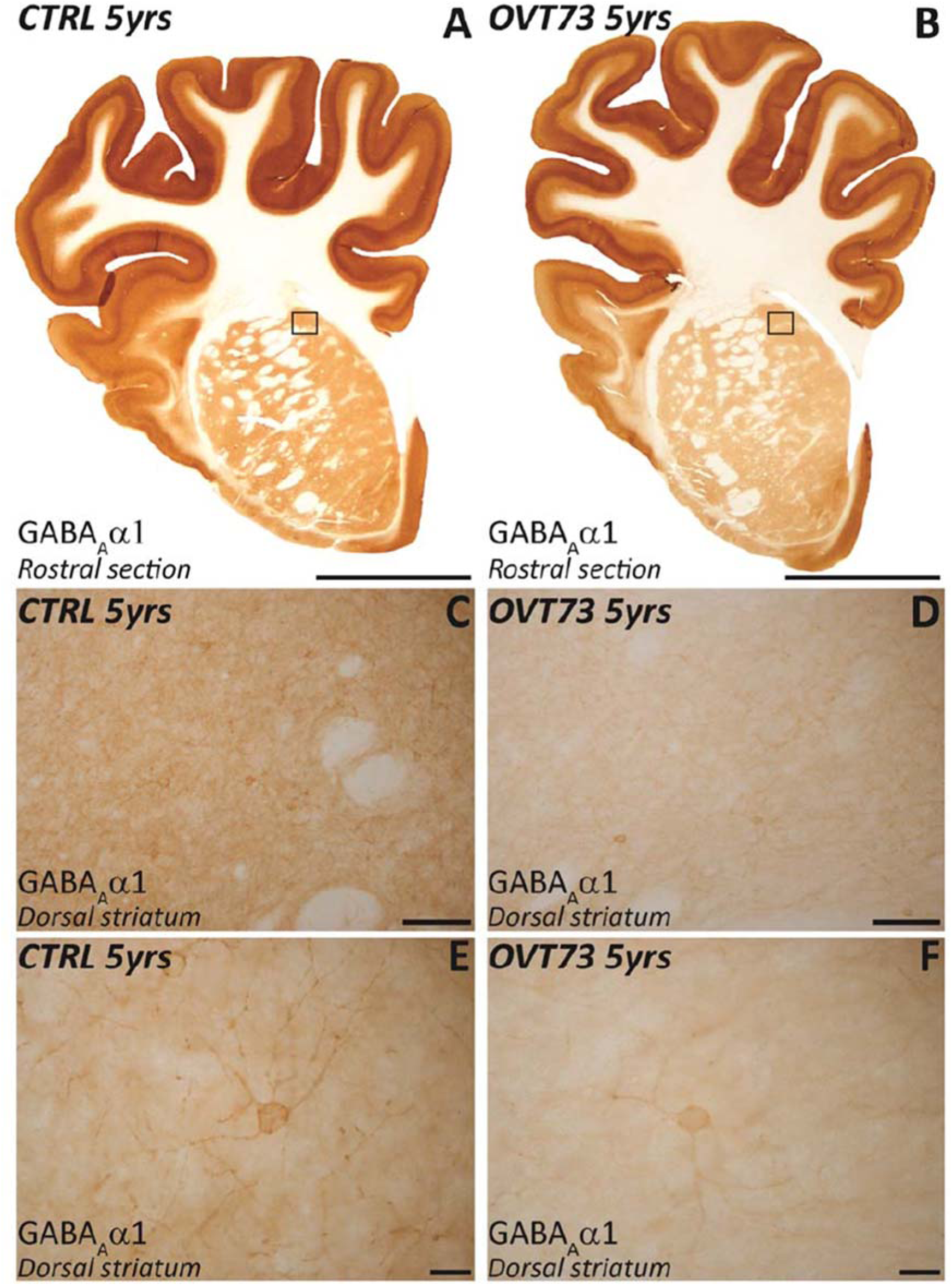
GABA_A_α1 density decrease in dorsal striatum of *OVT73* animals. These representative images show singleLlabel DAB immunohistochemistry for GABA_A_α1 in the striatum (rostral section) in 5-year-old control and *OVT73* animals. (A & B) Macros of the right hemispheres of the control and *OVT73* animals stained with GABA_A_α1 antibody. An overall reduced intensity of staining can be observed macroscopically in the dorsal striatum but not ventral striatum of the *OVT73* animals. (C & D) A clear reduction of GABA_A_α1 can be noticed at higher magnification (20x). (E & F) The high-resolution photomicrographs (63x) show a decrease of GABA_A_α1 receptor intensity across the perikarya and neuritic processes of GABAergic neurons. Scale bars in A and B = 1 cm, C and D = 100 μm, E and F = 20 μm.

**Figure 10.**
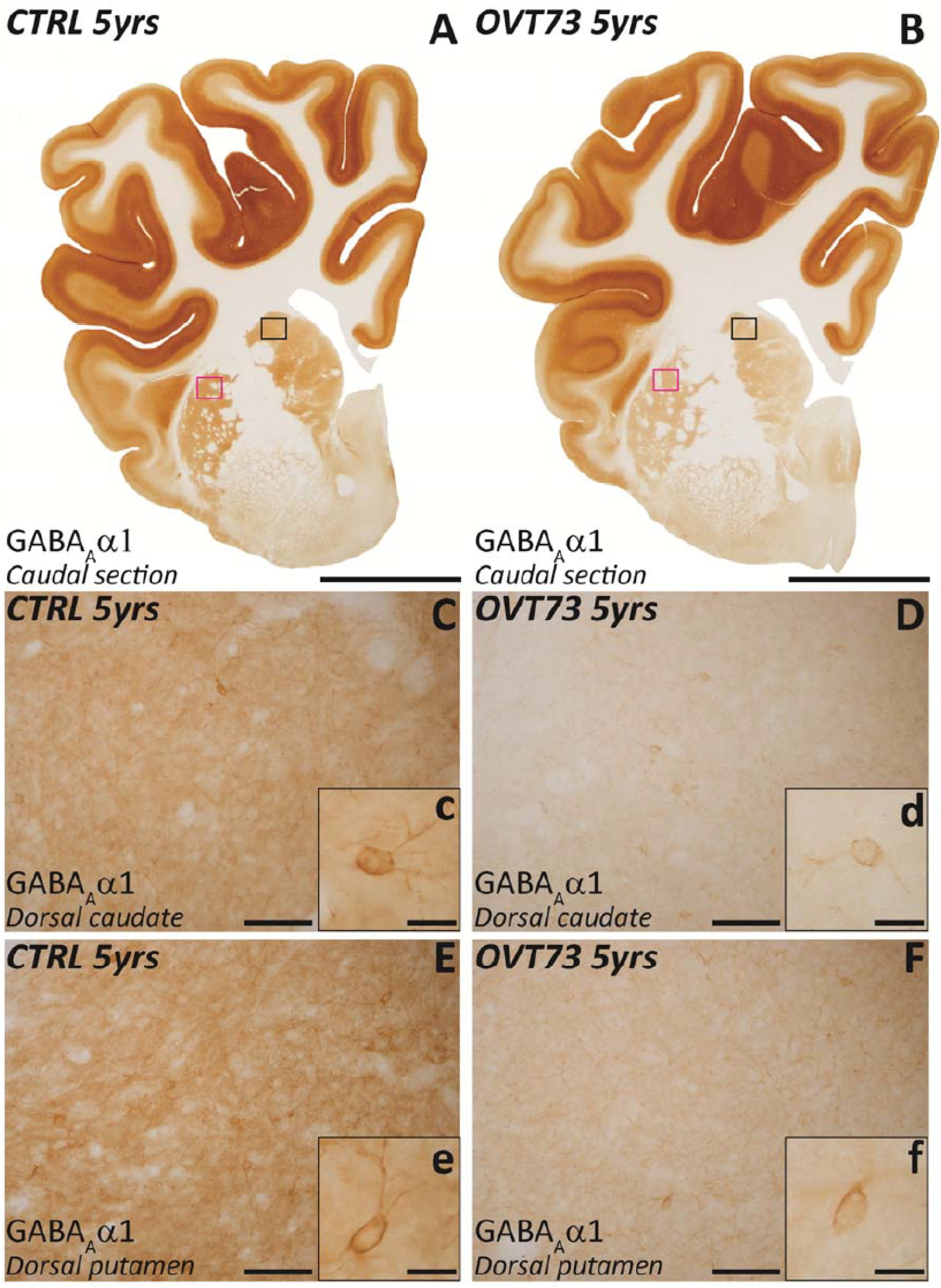
GABA_A_α1 density decrease in dorsal caudate and dorsal putamen of *OVT73* animals. These representative images show the immunostaining for GABA_A_α1 in caudate nucleus (caudal section) and putamen (caudal section) in 5-year-old controls and *OVT73*. (A & B) Low magnification images of the right hemispheres of the control and *OVT73* animals immunolabelled for GABA_A_α1 antibody. A decrease of intensity of staining is apparent in the caudate nucleus (black square) and putamen (fuchsia square) of *OVT73* animals. Higher magnification images (20x) highlight the reduction of GABA_A_α1 immunolabelling in the dorsal caudate (C & D) and dorsal putamen (E and F) of *OVT73* animals. High resolution photomicrographs (63x) illustrate the loss of GABAAα1 receptor in the soma and dendritic processes of GABAergic cells in the dorsal caudate (c and d) and dorsal putamen (e and f) of *OVT73*. Scale bars in A and B = 1 cm, C,D,E,F = 100 μm, c,d,e,f = 20 μm.

### Gene regulatory networks show reduced CREB regulon activity in *OVT73*

We assessed the transcription factor regulation of genes differentially expressed in *OVT73* through construction of gene regulatory networks using SCENIC. SCENIC performs cis regulatory motif analysis to find putative direct binding sites between co-expressed transcription factors and genes. These co-expressed transcription factor and genes are termed regulons [49]. The activity of each regulon in *OVT73* and control cell types (regulon activity) were scored using the AUCell algorithm which computes the enrichment of regulons as an area under the recovery curve across a ranking of all genes in a cell based on their normalised expression values. A high regulon activity indicates genes within the regulon are positively regulated by the transcription factor.

We focused on the regulon activity of the cAMP-responsive element-binding protein (CREB) family of transcription factors that has been shown to be activated following prolonged glutamate excitotoxicity [72, 73]. Regulon activity of CREB transcription factors, *CREB1*, *ATF2*, *ATF4*, *ATF7* was detected in medium spiny neurons, astrocytes, microglia, neuroblasts, oligodendrocytes and endothelial cells (Figure 11). When examining differential regulon activity between *OVT73* and controls, we observed a reduction in regulon activity for *CREB1* in astrocytes (differential module activity of -0.952, p = 0.0005, randomised permutation test with 2,000 permutations), D1 medium spiny neurons (−-1.43, p < 0.0005), D2 medium spiny neurons (−-1.367, p < 0.0005), eccentric medium spiny neurons (−-1.971, p < 0.0005), microglia (−-0.81, p < 0.0005), neuroblasts (−-0.83, p < 0.0005), oligodendrocytes (- 1.101, p = 0.0015) and OPC (−-0.894, p < 0.0005). *ATF2* regulon activity was reduced between *OVT73* and controls for D2 MSN (−-0.755, p < 0.0005), microglia (−-0.464, p < 0.0005), neuroblasts (−-0.555, p < 0.011), oligodendrocytes (−-0.364, p < 0.011), OPC (−-0.069, p < 0.0005). *ATF4* regulon activity was reduced between *OVT73* and controls for astrocytes (−-0.162, p = 0.001), D1 medium spiny neurons (−-0.087, p = 0.006), D2 medium spiny neurons (−-0.448, p < 0.0005), endothelial cells (−-1.541, p = 0.017), microglia (−-0.095, p = 0.0055), neuroblasts (−-0.250, p < 0.0005), oligodendrocytes (−-0.230, p < 0.0005), OPC (−-0.175, p < 0.0005) and SST/NPY interneurons (−-0.311, p = 0.0265). *ATF7* regulon activity was reduced between *OVT73* and controls for D2 medium spiny neurons (−-0.265, p < 0.0005), OPC (- 0.092, p < 0.0005) and PV/Th interneurons (−-0.214, p = 0.007) (Figure 11C, Supplementary Figure 13). The greatest sum of difference in CREB regulon activity between *OVT73* and controls (sum of *CREB1*, *ATF2*, *ATF4*, *ATF7* regulon activity in *OVT73* subtracted by the sum of *CREB1*, *ATF2*, *ATF4*, *ATF7* regulon activity in control) was observed in the D2 medium spiny neurons. Interestingly, CREB regulon activity was reduced to a lesser degree in D1 medium spiny neurons compared to D2 medium spiny neurons. Regulon activity of all identified regulons is available in Supplementary Figure 11 and Supplementary Figure 12. Gene members of each regulon are available in Supplementary File 8.

**Figure 11.**
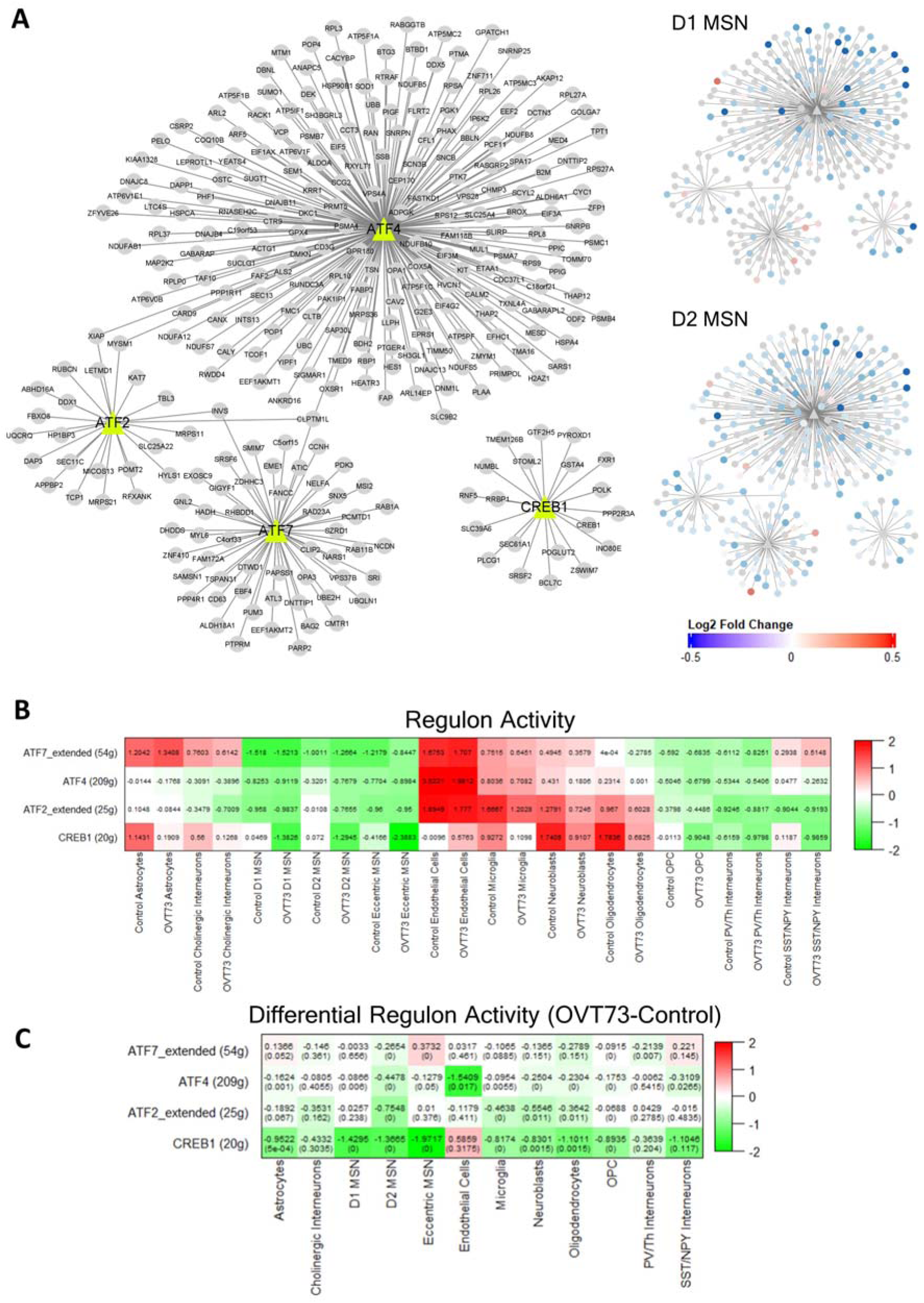
Gene regulatory networks show reduced CREB regulon activity in *OVT73*. (A) CREB-related transcription factor regulated gene modules (regulons) network. The associated transcription factor regulon is shown as triangles with gene members of the regulon connected as dots. The two panels on the right show the log2 fold change of differentially expressed gene members of the regulon between *OVT73* and controls in D1 medium spiny neurons and D2 medium spiny neurons. (B) Heatmap of CREB regulon activity in *OVT73* and control cell types. Regulon activity was computed based on transcription factor regulated gene modules using *OVT73* differentially expressed genes as input. (C) Differential regulon activity was computed by subtraction of *OVT73* regulon activity and control regulon activity. A randomised permutation test with 2,000 permutations was performed to determine significant differential regulon activity between OVT73 and control cell types. P-values of the randomised permutation test are shown in the parentheses.

## Discussion

### Evidence for glutamate excitotoxicity in the *OVT73* medium spiny neurons

Investigations of pathogenesis prior to the onset of striatal cell loss and motor and cognitive deficits is vital for the understanding of HD disease initiating events. Our group has generated and characterised a transgenic sheep model of HD (named *OVT73*) that displays no striatal cell loss or overt symptoms but exhibits many of the molecular changes reminiscent of the prodromal phase of human disease [19–24]. We set out to examine single nuclei transcriptomes of the cell types present in the *OVT73* striatum to investigate potential pathogenic events leading to the onset of cell death. Our single nuclei RNA-seq of the *OVT73* striatum reveals transcriptomic evidence to support increased synaptic signalling in the *OVT73* medium spiny neurons. The combination of differential gene expression analysis, gene co-expression network analysis and cell-cell signalling analysis indicated the glutamatergic and GABAergic synapse as an early site affected in the *OVT73* HD striatum.

With respect to the glutamatergic synapse, we observed an upregulation of genes encoding ionotropic glutamate receptors including NMDA, AMPA and kainate receptors in *OVT73* D1 and D2 medium spiny neurons compared to controls. Co-expression analysis revealed increased activity of the gene modules containing these glutamate receptor genes in *OVT73* D1 and D2 medium spiny neurons. Moreover, cell-cell signalling inferred from ligand receptor expression showed increased signalling for pathways including neurexin, neuroligin, neuregulin, neural cell adhesion molecule and ephrin. These signalling pathways have been implicated in neurite outgrowth, glutamatergic synapse formation and have been shown to increase ionotropic glutamate receptor abundance [59, 74–84]. Prior evidence indicates a positive correlation between ionotropic glutamate receptor mRNA levels, receptor protein abundance and receptor signalling inferred from ligand binding studies and changes in receptor currents [85–89]. Therefore, it is likely that an upregulation of transcription from these ionotropic glutamate receptor genes results in an increased abundance of these synaptic receptors and increased signalling.

The glutamate excitotoxicity hypothesis proposes neuronal stress leading to initiation of cell death pathways arising from excessive glutamate receptor signalling. Our observations provide evidence to support the activity of this excitotoxic process in the *OVT73* striatum. As further support, multiple studies investigating glutamate receptor binding and electrophysiological analyses of glutamate receptor currents have indicated that glutamate receptor signalling is increased during the presymptomatic phase in HD mouse models. Although the increase in glutamate signalling was transient and gradually decreased with disease progression and increasing cell loss with reduced numbers of glutamate receptors [27–29, 85, 90, 91]. Since the 5-year-old *OVT73* HD sheep exhibit no striatal cell loss, our detection of an upregulation of transcription for ionotropic glutamate receptors in *OVT73* D1 and D2 medium spiny neurons is likely reflective of an early disease phase in which neurons are still able to cope with the level of excitotoxic stress and is consistent with observed increased glutamate signalling during the early stages of HD.

The overactivation of ionotropic glutamate receptors leads to an influx of calcium ions promoting oxidative stress, mitochondrial dysfunction, and eventual initiation of cell death [7, 92]. Mitochondrial dysfunction and oxidative stress are common themes in HD evidenced by a decrease in the activity of oxidative phosphorylation (OXPHOS) complexes [93–95]. A recent single nuclei RNA-seq study of the HD striatum in human and HD mouse models also provides transcriptomic evidence for a downregulation of these OXPHOS complexes in the medium spiny neurons [43]. Similarly, we have detected a downregulation of transcription from genes encoding OXPHOS complexes including complex I, II, III and IV and V in the *OVT73* D1 and D2 medium spiny neurons. We postulate that the downregulation of these OXPHOS complexes is a direct consequence of the glutamate excitotoxicity process. The influx of calcium ions mediated by ionotropic glutamate receptors can also trigger neuroprotective pathways including phosphorylation of the transcription factor CREB and activation by CREB binding protein [72]. CREB plays a role in a wide range of neuroprotective processes including maintenance of synaptic plasticity, neurogenesis, neuronal growth, and survival [96, 97]. Studies have indicated that CREB function is lost in HD prior to cell death [73] and may be directly regulated by mutant huntingtin (m*HTT*) [98–100]. We have observed both a downregulation of genes encoding CREB transcription factors (*CREB1*, *ATF2*, *ATF4* and *ATF7*) and a reduction of CREB regulon activity inferred from gene regulatory networks across *OVT73* D1 and D2 medium spiny neurons, astrocytes, microglia, neuroblasts and oligodendrocytes. The greatest reduction in CREB regulon activity was identified in the *OVT73* D2 medium spiny neurons. Our results provide further evidence for CREB transcriptional dysregulation in HD that is potentially indicative of the initiating stages of a cell death cascade.

We have presented observations that support increased glutamate excitotoxicity vulnerability in *OVT73* D2 medium spiny neurons over D1 medium spiny neurons. Genes encoding NMDA receptor subunit NR2B and AMPA receptor subunits 3-5 were exclusively upregulated in D2 medium spiny neurons. Cell-cell signalling pathways including neurexin, neuregulin and ephrin that mediate glutamatergic and GABAergic synaptogenesis [59, 74–84] showed increased number of ligand-receptor interactions and higher probability of communication in D2 relative to D1 medium spiny neurons. Furthermore, gene regulatory network analysis revealed a stronger reduction in CREB transcription factor regulon activity in D2 medium spiny neurons compared to D1 medium spiny neurons. These observations are consistent with reports that D2 medium spiny neurons are preferentially lost in prodromal disease which may correlate to the clinical presentation of chorea symptoms early in HD [101, 102].

### Compensatory mechanisms that act to reduce excitotoxicity

In human HD, somatic instability has been observed in the striatum [103–107] with the glutamine coding tract becoming progressively expanded throughout disease course [108]. It is proposed that excitotoxic stress becomes progressively worse as the repeat undergoes somatic expansion until the cells can no longer cope and cell death processes begin [108]. As previously discussed, *OVT73* likely models a prodromal phase of HD. The 73-unit polyglutamine repeat coding tract is stable in somatic cells and combined with the observation that there is an absence of striatal neuronal loss, it is likely that there are compensatory mechanisms allowing the cells to cope with the level of excitotoxic stress. The main mechanism for synaptic glutamate removal in the brain is through uptake into astrocytes via glutamate transporters and degradation via the glutamine-glutamate cycle [109, 110]. We have observed an increase in the mRNA expression of *SLC1A2* and *SLC1A3* genes that encode the glutamate uptake transporters, GLT1 and GLAST respectively in *OVT73* astrocytes compared to control astrocytes which may represent a compensatory response. Gene co-expression network analysis also indicate an increased activity of modules containing the *SLC1A2* and *SLC1A3* genes in *OVT73* compared to control astrocytes. Further, cell-cell signalling inferred from ligand-receptor expression revealed an increase in EHPA and FGF signalling pathways which have been shown to promote the expression of *SLC1A2* and *SLC1A3* [81, 111, 112]. Previous studies have shown that *SLC1A2* and *SLC1A3* mRNA levels were positively correlated with GLT1 and GLAST mediated uptake of synaptic glutamate [89, 113, 114]. Moreover, experimentally induced upregulation of GLT1 transporter expression by ceftriaxone increased astrocytic synaptic glutamate uptake and ameliorated motor dysfunction in the R6/2 HD mouse model [115, 116]. We postulate that the upregulation of *SLC1A2* and *SLC1A3* in the *OVT73* sheep was in response, to remove excess synaptic glutamate. Studies in the HD striatum of post-mortem human brains and HD mouse models show diminished levels of GLT1/GLAST mRNA, protein, and reduced glutamate uptake into astrocytes. This reduction in glutamate uptake was hypothesised to exacerbate glutamate excitotoxicity [89, 114, 117–120]. It is likely that the upregulation of genes encoding glutamate uptake transporters observed in our study is reflective of an earlier disease stage before the onset of neurodegeneration and where multiple compensatory mechanisms are still active to combat excitotoxicity.

### Upregulation of GABA_A_ receptor genes in *OVT73* medium spiny neurons

We also observed a transcriptional upregulation of genes encoding GABA_A_ receptor subunits α, β, and γ in *OVT73* D1 and D2 medium spiny neurons compared to controls. In support of transcript levels representing protein levels, the mRNA levels of GABA_A_ receptors have been found to be positively correlated with receptor protein abundance and receptor signalling inferred through receptor binding studies and changes in receptor currents [121, 122]. Based on our transcript observations, we propose that increased GABA_A_ receptor signalling is likely occurring in the *OVT73* striatum. Striatal GABA_A_ currents have been postulated to have a neuroprotective effect against excitotoxicity [123, 124] and therefore increased GABA_A_ signalling in *OVT73* may be another mechanism that mitigates excitotoxic stress providing neuronal protection in the *OVT73* striatum. We have also described an upregulation of genes encoding GABA transporters (*SLC6A1*, *SLC6A11*) and the glutamate to GABA conversion gene, glutamate decarboxylase (*GAD1* and *GAD2*). Elevated levels of GABA have been associated with an increased transcription of GABA transporters [125] and glutamate decarboxylases [126–128]. From these observations, we propose that the increases in GABA_A_ receptor signalling may be a result of increased synaptic GABA concentrations resulting from an increased conversion of glutamate to GABA.

The biology surrounding GABA_A_ receptor mRNA and protein expression and receptor signalling in HD is complex, with differential regulation of various GABA_A_ receptor subunits that is dependent on disease stage [121, 122]. The consensus is that a reduction in GABA_A_ receptor binding and reduced GABA_A_ currents are present during the symptomatic disease phase of HD individuals and HD mouse models [121, 122]. Therefore, it is possible that the increased GABA_A_ receptor signalling identified in *OVT73* is yet another early disease phase event compensating for the glutamate excitotoxic stress and this neuroprotective effect is lost with disease progression. This is consistent with reports of increased GABA_A_ signalling in presymptomatic HD mouse models that is reduced during later disease stages [129, 130].

Despite an increase in GABA_A_α1 transcriptomic expression, our immunohistochemical densitometric analyses showed no difference in the immunoreactivity of the GABA_A_α1 receptor subunit between *OVT73* and controls in the ventral region of the striatum that comprised the tissue utilised for the single nuclei RNA-seq experiment. This discrepancy may be explained by a transcriptional fold change (fold change of 1.07) that is not at a level detectable by the immunohistochemical assay. Alternatively, transcriptional upregulation of GABA_A_α1 was observed only in a subset of cell types including *OVT73* D1, D2 medium spiny neurons, PV/Th interneurons and OPC and therefore this increase in expression may not be observable in the immunohistochemical assay that is averaged over all cell types. Post transcriptional and post-translational regulation of GABA_A_α1 could also explain the difference. We have however described a 35%-39% decrease in GABA_A_α1 immunoreactivity in the dorsal striatal regions. This is an intriguing observation and suggests regional variation in receptor expression. Magnetic resonance imaging (MRI) studies have shown that dorsal and medial regions of the HD striatum exhibit greater atrophy than ventral regions [131–133]. Our observations of a decrease in GABA_A_α1 receptor immunoreactivity in dorsal regions may provide the first indications of an overwhelming of the GABA_A_ signalling pathways in the dorsal regions which are preferentially affected early in the disease. A potential regional excitotoxicity effect may be responsible for the overwhelming of GABA signalling pathways although further studies will need to be performed to address this effect.

### Limitations

A limitation of our study is the small sample size comprising six *OVT73* and six controls. This may restrict our ability to detect statistically significant effects of minor magnitude. However, this cohort of animals has been extensively characterised by transcriptomics [25], proteomics [25], metabolomics [20, 21, 24], and behavioural analyses [22, 23]. These studies have successfully captured many features of prodromal HD. Moreover, the striatal tissues utilised in this study were from the same region used in a bulk RNA-seq study that identified transcriptomic differences [25]. All the datasets from the previous studies of this cohort of animals are available on a publicly accessible database [26]. Additionally, we also recognise the necessity for functional analyses to confirm the transcriptional differences identified in the present data. These may include characterisation of glutamate and GABA receptor binding, electrophysiological assessments of glutamate and GABA currents, and validation at the protein level using immunohistochemistry methods. Unfortunately, striatal tissue availability is scarce, and tissue resources from many of these animals are depleted. Future functional characterisation in progeny cohorts will be critical, not only to confirm results summarised in this study, but also for comprehensive characterisation of the overall *OVT73* sheep model.

### The case for early therapeutic intervention

The detection of an excitotoxic process in the *OVT73* striatum with minimal pathology and no striatal cell loss suggests that excitotoxic stress may be detected long before the initiation of the cell death cascade. The observations in our single nuclei RNA-seq of the prodromal *OVT73* model corroborates reports in HD mouse models that showcase excitotoxicity begins early on in disease phase before the onset of symptoms [27–29]. Current therapeutics targeting excitotoxicity have included NMDA receptor antagonists (Memantine, Amantadine) that reduce the overactivation of NMDA receptors and these have been utilised in clinical trials to limited success [134–137]. One potential explanation for the lack of efficacy is that the therapeutic agents were administrated too late in disease stage. A majority of HD clinical trials have eligibility criteria’s that require the potential participants to exhibit reduced motor or cognitive function (i.e., reduction in UHDRS total functional capacity). However, at this stage, widespread cellular atrophy would have already occurred in the striatum and other brain regions. Disease treatments may be confounded by the consequences of widespread catastrophe brought upon by the cell death cascade reaching a point where recovery becomes unattainable. Further, the cell death cascade releases numerous neuroinflammatory cytokines, proinflammatory markers and reactive oxygen species that can affect a wide range of biological pathways complicating the initial targets derived from the therapeutic agent [138]. An example was seen in Alzheimer’s disease where the accumulation of pathology in amyloid plaques and tau tangles over many years rendered treatments that act to reduce these pathogenic species ineffective [139, 140]. Similarly, this may be the case in HD, where accumulation of cell death could result in poor translation of therapeutics that act to reduce excitotoxicity. In support of this, striatal neurons of HD mouse models have shown to gradually become resistant to excitotoxicity modifying agents following disease progression [141–144]. The findings of our single nuclei RNA-seq study advocates for early intervention before the onset of the cell death cascade which may lead to different clinical outcomes for many of tested therapeutic agents.

## Conclusions

In summary, our study of the single nuclei transcriptome of a Huntington’s disease prodromal model has provided robust evidence for glutamate excitotoxicity in the HD striatum occurring before the onset of neurodegeneration and motor symptoms (overview of glutamate excitotoxicity in the *OVT73* model shown in Figure 12). Moreover, we have revealed that medium spiny neurons the most vulnerable cell type in HD, were specifically found to have differentially expressed transcriptional markers of a response to the proposed consequences of a glutamate excitotoxic process. We have additionally identified reduced transcriptional activity of CREB, thought to be a downstream effector of excitotoxicity. Transcription from several potentially compensatory mechanisms that act to reduce excitotoxicity were detected in *OVT73* nuclei, including increased glutamate uptake into astrocytes, and increased GABAergic signalling. As we and others [27–29] have suggested the process of excitotoxicity occurs early in disease progression, early intervention before the onset of the cell death cascade could be critical. Taken together, the molecular phenotypes presented here position *OVT73* HD model as an excellent resource for mechanistically capturing the prodromal phase of the disease and for testing therapeutics targeted to address the condition at the presymptomatic phase.

**Figure 12.**
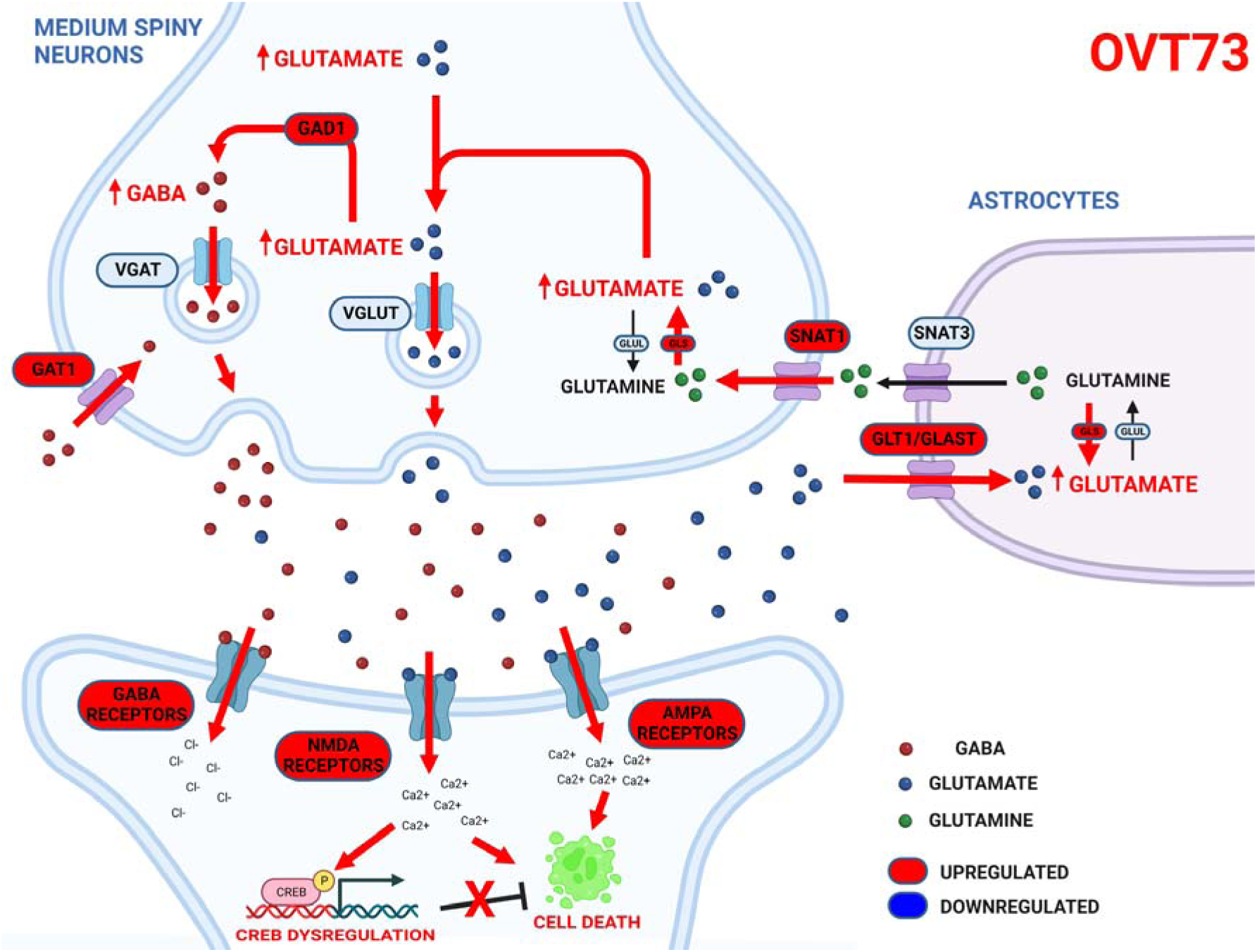
Proposed overview of glutamate excitotoxicity in the HD *OVT73* striatum. In the *OVT73* striatum, an excess of glutamate triggers overactivation of post-synaptic NDMA and AMPA receptors resulting in an influx of calcium ions and activation of cell death pathways. Anti-cell death pathways including CREB activation are dysregulated in *OVT73*. Synaptic glutamate is transported into astrocytes by GLT1/GLAST transporters acting to decrease levels of synaptic glutamate and potential allowing the cells to cope with the level of excitotoxic stress. GABA synthesis from glutamate is increased and results in increased GABA receptor signalling. Created with BioRender.com.

## Ethics approval and consent to participate

All experiments were performed in accordance with the SARDI/PIRSA Animal Ethics Committee (approval no. 19/02 and 05/12). All experiments performed adhered to the recommendations in the ARRIVE guidelines [30]. No patients or human data was used in this study.

## Consent for publication

All authors have read the manuscript and approved to be co-authors on the manuscript.

## Availability of data and materials

The data have been deposited in NCBI’s Gene Expression Omnibus (GEO) [145] and are accessible through accession number GSE229839 (https://www.ncbi.nlm.nih.gov/geo/query/acc.cgi?acc= GSE229839).

The code used in this study are deposited on zenodo available at https://doi.org/10.5281/zenodo.8057929.

## Authors’ contributions

A.J., L.Y., R.R.H., J.F.G., M.E.M., H.J.W., R.L.M.F., K.L. and R.G.S. considered the study. A.J. and V.H. performed the single nuclei RNA-seq library preparation. R.R.H., S.J.R., J.C.J., S.R.R., C.J.M., J.M.K., harvested the striatal tissue. S.P. performed the immunohistochemistry experiments. S.R.R., C.J.M., J.M.K., P.J.V. and C.S.B was responsible for the on-farm management of the *OVT73* animals. A.J. and L.Y. performed the single nuclei RNA-seq analysis and interpretation. A.J. wrote the manuscript. All the authors read and approved the final manuscript.

## Acknowledgements

The authors wish to thank the incredible on-farm SARDI team for all-animal management. Additionally, we thank the New Zealand–China Non-Communicable Diseases Research Collaboration Centre (NCD CRCC).

## References

1. The Huntington’s Disease Collaborative Research Group: A novel gene containing a trinucleotide repeat that is expanded and unstable on Huntington’s disease chromosomes. The Huntington’s Disease Collaborative Research Group. Cell 1993, 72(6):971–983.

2. Snell RG, MacMillan JC, Cheadle JP, Fenton I, Lazarou LP, Davies P, MacDonald ME, Gusella JF, Harper PS, Shaw DJ: Relationship between trinucleotide repeat expansion and phenotypic variation in Huntington’s disease. Nat Genet 1993, 4(4):393–397.

3. Doble A: The role of excitotoxicity in neurodegenerative disease: implications for therapy. Pharmacol Ther 1999, 81(3):163–221.

4. Salinska E, Danysz W, Lazarewicz JW: The role of excitotoxicity in neurodegeneration. Folia Neuropathol 2005, 43(4):322–339.

5. Dong X-x, Wang Y, Qin Z-h: Molecular mechanisms of excitotoxicity and their relevance to pathogenesis of neurodegenerative diseases. Acta Pharmacologica Sinica 2009, 30(4):379–387.

6. Mehta A, Prabhakar M, Kumar P, Deshmukh R, Sharma PL: Excitotoxicity: Bridge to various triggers in neurodegenerative disorders. European Journal of Pharmacology 2013, 698(1):6–18.

7. Sepers MD, Raymond LA: Mechanisms of synaptic dysfunction and excitotoxicity in Huntington’s disease. Drug Discov Today 2014, 19(7):990–996.

8. Beal MF, Kowall NW, Ellison DW, Mazurek MF, Swartz KJ, Martin JB: Replication of the neurochemical characteristics of Huntington’s disease by quinolinic acid. Nature 1986, 321(6066):168–171.

9. Bordelon YM, Chesselet M-F, Nelson D, Welsh F, Erecińska M: Energetic Dysfunction in Quinolinic Acid-Lesioned Rat Striatum. Journal of Neurochemistry 1997, 69(4):1629–1639.

10. Emerich DF, Thanos CG, Goddard M, Skinner SJ, Geany MS, Bell WJ, Bintz B, Schneider P, Chu Y, Babu RS et al: Extensive neuroprotection by choroid plexus transplants in excitotoxin lesioned monkeys. Neurobiol Dis 2006, 23(2):471–480.

11. Foster AC, Miller LP, Oldendorf WH, Schwarcz R: Studies on the disposition of quinolinic acid after intracerebral or systemic administration in the rat. Exp Neurol 1984, 84(2):428–440.

12. Beal MF, Ferrante RJ, Swartz KJ, Kowall NW: Chronic quinolinic acid lesions in rats closely resemble Huntington’s disease. J Neurosci 1991, 11(6):1649–1659.

13. Coyle JT, Schwarcz R: Lesion of striatal neurones with kainic acid provides a model for Huntington’s chorea. Nature 1976, 263(5574):244–246.

14. McGeer EG, McGeer PL: Duplication of biochemical changes of Huntington’s chorea by intrastriatal injections of glutamic and kainic acids. Nature 1976, 263(5577):517–519.

15. Milnerwood AJ, Gladding CM, Pouladi MA, Kaufman AM, Hines RM, Boyd JD, Ko RW, Vasuta OC, Graham RK, Hayden MR et al: Early increase in extrasynaptic NMDA receptor signaling and expression contributes to phenotype onset in Huntington’s disease mice. Neuron 2010, 65(2):178–190.

16. Joshi PR, Wu NP, Andre VM, Cummings DM, Cepeda C, Joyce JA, Carroll JB, Leavitt BR, Hayden MR, Levine MS et al: Age-dependent alterations of corticostriatal activity in the YAC128 mouse model of Huntington disease. J Neurosci 2009, 29(8):2414–2427.

17. Heng MY, Detloff PJ, Wang PL, Tsien JZ, Albin RL: In vivo evidence for NMDA receptor mediated excitotoxicity in a murine genetic model of Huntington disease. J Neurosci 2009, 29(10):3200–3205.

18. Jacobsen JC, Bawden CS, Rudiger SR, McLaughlan CJ, Reid SJ, Waldvogel HJ, MacDonald ME, Gusella JF, Walker SK, Kelly JM et al: An ovine transgenic Huntington’s disease model. Hum Mol Genet 2010, 19(10):1873–1882.

19. Huntington’s Disease Sheep Collaborative Research G, Reid SJ, Patassini S, Handley RR, Rudiger SR, McLaughlan CJ, Osmand A, Jacobsen JC, Morton AJ, Weiss A et al: Further molecular characterisation of the OVT73 transgenic sheep model of Huntington’s disease identifies cortical aggregates. J Huntingtons Dis 2013, 2(3):279–295.

20. Handley RR, Reid SJ, Patassini S, Rudiger SR, Obolonkin V, McLaughlan CJ, Jacobsen JC, Gusella JF, MacDonald ME, Waldvogel HJ et al: Metabolic disruption identified in the Huntington’s disease transgenic sheep model. Sci Rep 2016, 6:20681.

21. Skene DJ, Middleton B, Fraser CK, Pennings JLA, Kuchel TR, Rudiger SR, Bawden CS, Morton AJ: Metabolic profiling of presymptomatic Huntington’s disease sheep reveals novel biomarkers. Scientific Reports 2017, 7(1):43030.

22. Morton AJ, Rudiger SR, Wood NI, Sawiak SJ, Brown GC, McLaughlan CJ, Kuchel TR, Snell RG, Faull RL, Bawden CS: Early and progressive circadian abnormalities in Huntington’s disease sheep are unmasked by social environment. Hum Mol Genet 2014, 23(13):3375–3383.

23. Vas S, Nicol AU, Kalmar L, Miles J, Morton AJ: Abnormal patterns of sleep and EEG power distribution during non-rapid eye movement sleep in the sheep model of Huntington’s disease. Neurobiology of Disease 2021, 155:105367.

24. Morton AJ, Middleton B, Rudiger S, Bawden CS, Kuchel TR, Skene DJ: Increased plasma melatonin in presymptomatic Huntington disease sheep (Ovis aries): Compensatory neuroprotection in a neurodegenerative disease? Journal of Pineal Research 2020, 68(2):e12624.

25. Handley RR, Reid SJ, Brauning R, Maclean P, Mears ER, Fourie I, Patassini S, Cooper GJS, Rudiger SR, McLaughlan CJ et al: Brain urea increase is an early Huntington’s disease pathogenic event observed in a prodromal transgenic sheep model and HD cases. Proc Natl Acad Sci U S A 2017, 114(52):E11293–E11302.

26. Mears ER, Handley RR, Grant MJ, Reid SJ, Day BT, Rudiger SR, McLaughlan CJ, Verma PJ, Bawden SC, Patassini S et al: A Multi-Omic Huntington’s Disease Transgenic Sheep-Model Database for Investigating Disease Pathogenesis. J Huntingtons Dis 2021, 10(4):423–434.

27. Levine MS, Klapstein GJ, Koppel A, Gruen E, Cepeda C, Vargas ME, Jokel ES, Carpenter EM, Zanjani H, Hurst RS et al: Enhanced sensitivity to N-methyl-D-aspartate receptor activation in transgenic and knockin mouse models of Huntington’s disease. J Neurosci Res 1999, 58(4):515–532.

28. Cepeda C, Ariano MA, Calvert CR, Flores-Hernandez J, Chandler SH, Leavitt BR, Hayden MR, Levine MS: NMDA receptor function in mouse models of Huntington disease. J Neurosci Res 2001, 66(4):525–539.

29. Starling AJ, Andre VM, Cepeda C, de Lima M, Chandler SH, Levine MS: Alterations in N-methyl-D-aspartate receptor sensitivity and magnesium blockade occur early in development in the R6/2 mouse model of Huntington’s disease. J Neurosci Res 2005, 82(3):377–386.

30. Percie du Sert N, Hurst V, Ahluwalia A, Alam S, Avey MT, Baker M, Browne WJ, Clark A, Cuthill IC, Dirnagl U et al: The ARRIVE guidelines 2.0: Updated guidelines for reporting animal research. PLOS Biology 2020, 18(7):e3000410.

31. Waldvogel HJ, Kubota Y, Fritschy J, Mohler H, Faull RL: Regional and cellular localisation of GABA(A) receptor subunits in the human basal ganglia: An autoradiographic and immunohistochemical study. J Comp Neurol 1999, 415(3):313–340.

32. Krishnaswami SR, Grindberg RV, Novotny M, Venepally P, Lacar B, Bhutani K, Linker SB, Pham S, Erwin JA, Miller JA et al: Using single nuclei for RNA-seq to capture the transcriptome of postmortem neurons. Nature Protocols 2016, 11(3):499–524.

33. Liao Y, Smyth GK, Shi W: featureCounts: an efficient general purpose program for assigning sequence reads to genomic features. Bioinformatics 2014, 30(7):923–930.

34. Liao Y, Smyth GK, Shi W: The Subread aligner: fast, accurate and scalable read mapping by seed-and-vote. Nucleic Acids Res 2013, 41(10):e108.

35. Danecek P, Bonfield JK, Liddle J, Marshall J, Ohan V, Pollard MO, Whitwham A, Keane T, McCarthy SA, Davies RM et al: Twelve years of SAMtools and BCFtools. Gigascience 2021, 10(2).

36. Xu J, Falconer C, Nguyen Q, Crawford J, McKinnon BD, Mortlock S, Senabouth A, Andersen S, Chiu HS, Jiang L et al: Genotype-free demultiplexing of pooled single-cell RNA-seq. Genome Biology 2019, 20(1):290.

37. Jiang A, Lehnert K, You L, Snell RG: ICARUS, an interactive web server for single cell RNA-seq analysis. Nucleic Acids Research 2022, 50(W1):W427–W433.

38. Jiang A, You L, Snell RG, Lehnert K: Delineation of complex gene expression patterns in single cell RNA-seq data with ICARUS v2.0. NAR Genom Bioinform 2023, 5(2):lqad032.

39. Hao Y, Hao S, Andersen-Nissen E, Mauck WM, 3rd, Zheng S, Butler A, Lee MJ, Wilk AJ, Darby C, Zager M et al: Integrated analysis of multimodal single-cell data. Cell 2021, 184(13):3573–3587 e3529.

40. Korsunsky I, Millard N, Fan J, Slowikowski K, Zhang F, Wei K, Baglaenko Y, Brenner M, Loh P-r, Raychaudhuri S: Fast, sensitive and accurate integration of single-cell data with Harmony. Nature Methods 2019, 16(12):1289–1296.

41. Malaiya S, Cortes-Gutierrez M, Herb BR, Coffey SR, Legg SRW, Cantle JP, Colantuoni C, Carroll JB, Ament SA: Single-Nucleus RNA-Seq Reveals Dysregulation of Striatal Cell Identity Due to Huntington’s Disease Mutations. J Neurosci 2021, 41(25):5534–5552.

42. Saunders A, Macosko EZ, Wysoker A, Goldman M, Krienen FM, de Rivera H, Bien E, Baum M, Bortolin L, Wang S et al: Molecular Diversity and Specializations among the Cells of the Adult Mouse Brain. Cell 2018, 174(4):1015–1030.e1016.

43. Lee H, Fenster RJ, Pineda SS, Gibbs WS, Mohammadi S, Davila-Velderrain J, Garcia FJ, Therrien M, Novis HS, Gao F et al: Cell Type-Specific Transcriptomics Reveals that Mutant Huntingtin Leads to Mitochondrial RNA Release and Neuronal Innate Immune Activation. Neuron 2020, 107(5):891–908 e898.

44. Wu T, Hu E, Xu S, Chen M, Guo P, Dai Z, Feng T, Zhou L, Tang W, Zhan L et al: clusterProfiler 4.0: A universal enrichment tool for interpreting omics data. Innovation (N Y) 2021, 2(3):100141.

45. Morgan M, Shepherd L: AnnotationHub: client to access AnnotationHub resources. R package version 3.6.0. In.; 2022.

46. Song W-M, Zhang B: Multiscale Embedded Gene Co-expression Network Analysis. PLOS Computational Biology 2015, 11(11):e1004574.

47. Shannon P, Markiel A, Ozier O, Baliga NS, Wang JT, Ramage D, Amin N, Schwikowski B, Ideker T: Cytoscape: a software environment for integrated models of biomolecular interaction networks. Genome Res 2003, 13(11):2498–2504.

48. Yu G, Li F, Qin Y, Bo X, Wu Y, Wang S: GOSemSim: an R package for measuring semantic similarity among GO terms and gene products. Bioinformatics 2010, 26(7):976–978.

49. Aibar S, González-Blas CB, Moerman T, Huynh-Thu VA, Imrichova H, Hulselmans G, Rambow F, Marine J-C, Geurts P, Aerts J et al: SCENIC: single-cell regulatory network inference and clustering. Nature Methods 2017, 14(11):1083–1086.

50. Lambert SA, Jolma A, Campitelli LF, Das PK, Yin Y, Albu M, Chen X, Taipale J, Hughes TR, Weirauch MT: The Human Transcription Factors. Cell 2018, 175(2):598–599.

51. Huynh-Thu VA, Irrthum A, Wehenkel L, Geurts P: Inferring Regulatory Networks from Expression Data Using Tree-Based Methods. PLOS ONE 2010, 5(9):e12776.

52. Jin S, Guerrero-Juarez CF, Zhang L, Chang I, Ramos R, Kuan CH, Myung P, Plikus MV, Nie Q: Inference and analysis of cell-cell communication using CellChat. Nat Commun 2021, 12(1):1088.

53. Munoz-Manchado AB, Bengtsson Gonzales C, Zeisel A, Munguba H, Bekkouche B, Skene NG, Lonnerberg P, Ryge J, Harris KD, Linnarsson S et al: Diversity of Interneurons in the Dorsal Striatum Revealed by Single-Cell RNA Sequencing and PatchSeq. Cell Rep 2018, 24(8):2179–2190 e2177.

54. Aaronson J, Beaumont V, Blevins RA, Andreeva V, Murasheva I, Shneyderman A, Armah K, Gill R, Chen J, Rosinski J et al: HDinHD: A Rich Data Portal for Huntington’s Disease Research. J Huntingtons Dis 2021, 10(3):405–412.

55. Nelis E, Haites N, Van Broeckhoven C: Mutations in the peripheral myelin genes and associated genes in inherited peripheral neuropathies. Human Mutation 1999, 13(1):11–28.

56. Shy ME, Jáni A, Krajewski K, Grandis M, Lewis RA, Li J, Shy RR, Balsamo J, Lilien J, Garbern JY et al: Phenotypic clustering in MPZ mutations. Brain 2004, 127(2):371–384.

57. Gregorio I, Braghetta P, Bonaldo P, Cescon M: Collagen VI in healthy and diseased nervous system. Dis Model Mech 2018, 11(6).

58. Klimaschewski L, Claus P: Fibroblast Growth Factor Signalling in the Diseased Nervous System. Mol Neurobiol 2021, 58(8):3884–3902.

59. Craig AM, Kang Y: Neurexin-neuroligin signaling in synapse development. Curr Opin Neurobiol 2007, 17(1):43–52.

60. Mo X, Liu M, Gong J, Mei Y, Chen H, Mo H, Yang X, Li J: PTPRM Is Critical for Synapse Formation Regulated by Zinc Ion. Front Mol Neurosci 2022, 15:822458.

61. Chatterjee M, Schild D, Teunissen CE: Contactins in the central nervous system: role in health and disease. Neural Regen Res 2019, 14(2):206–216.

62. Luckenbill-Edds L: Laminin and the mechanism of neuronal outgrowth. Brain Res Brain Res Rev 1997, 23(1-2):1-27.

63. Jin J, Liu L, Chen W, Gao Q, Li H, Wang Y, Qian Q: The Implicated Roles of Cell Adhesion Molecule 1 (CADM1) Gene and Altered Prefrontal Neuronal Activity in Attention Deficit/Hyperactivity Disorder: A “Gene-Brain-Behavior Relationship”? Front Genet 2019, 10:882.

64. Miao H, Wang B: EphA receptor signaling--complexity and emerging themes. Semin Cell Dev Biol 2012, 23(1):16–25.

65. Gamez B, Rodriguez-Carballo E, Ventura F: BMP signaling in telencephalic neural cell specification and maturation. Front Cell Neurosci 2013, 7:87.

66. González-Castillo C, Ortuño-Sahagún D, Guzmán-Brambila C, Pallàs M, Rojas-Mayorquín AE: Pleiotrophin as a central nervous system neuromodulator, evidences from the hippocampus. Frontiers in Cellular Neuroscience 2015, 8.

67. Ditlevsen DK, Povlsen GK, Berezin V, Bock E: NCAM-induced intracellular signaling revisited. J Neurosci Res 2008, 86(4):727–743.

68. Ebnet K: Junctional Adhesion Molecules (JAMs): Cell Adhesion Receptors With Pleiotropic Functions in Cell Physiology and Development. Physiol Rev 2017, 97(4):1529–1554.

69. Bondy SC, LeBel CP: The relationship between excitotoxicity and oxidative stress in the central nervous system. Free Radic Biol Med 1993, 14(6):633–642.

70. Wullner U, Young AB, Penney JB, Beal MF: 3-Nitropropionic acid toxicity in the striatum. J Neurochem 1994, 63(5):1772–1781.

71. Kumar A, Ratan RR: Oxidative Stress and Huntington’s Disease: The Good, The Bad, and The Ugly. J Huntingtons Dis 2016, 5(3):217–237.

72. West AE, Chen WG, Dalva MB, Dolmetsch RE, Kornhauser JM, Shaywitz AJ, Takasu MA, Tao X, Greenberg ME: Calcium regulation of neuronal gene expression. Proceedings of the National Academy of Sciences 2001, 98(20):11024–11031.

73. Choi YS, Lee B, Cho HY, Reyes IB, Pu XA, Saido TC, Hoyt KR, Obrietan K: CREB is a key regulator of striatal vulnerability in chemical and genetic models of Huntington’s disease. Neurobiol Dis 2009, 36(2):259–268.

74. Budreck EC, Kwon O-B, Jung JH, Baudouin S, Thommen A, Kim H-S, Fukazawa Y, Harada H, Tabuchi K, Shigemoto R et al: Neuroligin-1 controls synaptic abundance of NMDA-type glutamate receptors through extracellular coupling. Proceedings of the National Academy of Sciences 2013, 110(2):725–730.

75. Espinosa F, Xuan Z, Liu S, Powell CM: Neuroligin 1 modulates striatal glutamatergic neurotransmission in a pathway and NMDAR subunit-specific manner. Frontiers in Synaptic Neuroscience 2015, 7.

76. Graf ER, Zhang X, Jin SX, Linhoff MW, Craig AM: Neurexins induce differentiation of GABA and glutamate postsynaptic specializations via neuroligins. Cell 2004, 119(7):1013–1026.

77. Sudhof TC: Neuroligins and neurexins link synaptic function to cognitive disease. Nature 2008, 455(7215):903-911.

78. Deng W, Luo F, Li B-m, Mei L: NRG1–ErbB4 signaling promotes functional recovery in a murine model of traumatic brain injury via regulation of GABA release. Experimental Brain Research 2019, 237(12):3351–3362.

79. Li B, Woo R-S, Mei L, Malinow R: The Neuregulin-1 Receptor ErbB4 Controls Glutamatergic Synapse Maturation and Plasticity. Neuron 2007, 54(4):583–597.

80. Noll JM, Li Y, Distel TJ, Ford GD, Ford BD: Neuroprotection by Exogenous and Endogenous Neuregulin-1 in Mouse Models of Focal Ischemic Stroke. Journal of Molecular Neuroscience 2019, 69(2):333–342.

81. Filosa A, Paixão S, Honsek SD, Carmona MA, Becker L, Feddersen B, Gaitanos L, Rudhard Y, Schoepfer R, Klopstock T et al: Neuron-glia communication via EphA4/ephrin-A3 modulates LTP through glial glutamate transport. Nature Neuroscience 2009, 12(10):1285–1292.

82. Dalva MB, Takasu MA, Lin MZ, Shamah SM, Hu L, Gale NW, Greenberg ME: EphB Receptors Interact with NMDA Receptors and Regulate Excitatory Synapse Formation. Cell 2000, 103(6):945–956.

83. Fux CM, Krug M, Dityatev A, Schuster T, Schachner M: NCAM180 and glutamate receptor subtypes in potentiated spine synapses: an immunogold electron microscopic study. Molecular and Cellular Neuroscience 2003, 24(4):939–950.

84. Sytnyk V, Leshchyns’ka I, Nikonenko AG, Schachner M: NCAM promotes assembly and activity-dependent remodeling of the postsynaptic signaling complex. Journal of Cell Biology 2006, 174(7):1071–1085.

85. Cha JH, Kosinski CM, Kerner JA, Alsdorf SA, Mangiarini L, Davies SW, Penney JB, Bates GP, Young AB: Altered brain neurotransmitter receptors in transgenic mice expressing a portion of an abnormal human huntington disease gene. Proc Natl Acad Sci U S A 1998, 95(11):6480–6485.

86. Cha JH, Frey AS, Alsdorf SA, Kerner JA, Kosinski CM, Mangiarini L, Penney JB, Jr., Davies SW, Bates GP, Young AB: Altered neurotransmitter receptor expression in transgenic mouse models of Huntington’s disease. Philos Trans R Soc Lond B Biol Sci 1999, 354(1386):981–989.

87. Landwehrmeyer GB, Standaert DG, Testa CM, Penney JB, Jr., Young AB: NMDA receptor subunit mRNA expression by projection neurons and interneurons in rat striatum. J Neurosci 1995, 15(7 Pt 2):5297–5307.

88. Ali NJ, Levine MS: Changes in expression of N-methyl-D-aspartate receptor subunits occur early in the R6/2 mouse model of Huntington’s disease. Dev Neurosci 2006, 28(3):230–238.

89. Arzberger T, Krampfl K, Leimgruber S, Weindl A: Changes of NMDA receptor subunit (NR1, NR2B) and glutamate transporter (GLT1) mRNA expression in Huntington’s disease-- an in situ hybridization study. J Neuropathol Exp Neurol 1997, 56(4):440–454.

90. Li L, Fan M, Icton CD, Chen N, Leavitt BR, Hayden MR, Murphy TH, Raymond LA: Role of NR2B-type NMDA receptors in selective neurodegeneration in Huntington disease. Neurobiol Aging 2003, 24(8):1113–1121.

91. Li L, Murphy TH, Hayden MR, Raymond LA: Enhanced striatal NR2B-containing N-methyl-D-aspartate receptor-mediated synaptic currents in a mouse model of Huntington disease. J Neurophysiol 2004, 92(5):2738–2746.

92. Pregi N, Belluscio LM, Berardino BG, Castillo DS, Canepa ET: Oxidative stress-induced CREB upregulation promotes DNA damage repair prior to neuronal cell death protection. Mol Cell Biochem 2017, 425(1-2):9–24.

93. Browne SE, Bowling AC, MacGarvey U, Baik MJ, Berger SC, Muqit MM, Bird ED, Beal MF: Oxidative damage and metabolic dysfunction in Huntington’s disease: selective vulnerability of the basal ganglia. Ann Neurol 1997, 41(5):646–653.

94. Brennan WA, Jr., Bird ED, Aprille JR: Regional mitochondrial respiratory activity in Huntington’s disease brain. J Neurochem 1985, 44(6):1948–1950.

95. Gu M, Gash MT, Mann VM, Javoy-Agid F, Cooper JM, Schapira AH: Mitochondrial defect in Huntington’s disease caudate nucleus. Ann Neurol 1996, 39(3):385–389.

96. Mantamadiotis T, Lemberger T, Bleckmann SC, Kern H, Kretz O, Villalba AM, Tronche F, Kellendonk C, Gau D, Kapfhammer J et al: Disruption of CREB function in brain leads to neurodegeneration. Nature Genetics 2002, 31(1):47–54.

97. Saura CA, Valero J: The role of CREB signaling in Alzheimer’s disease and other cognitive disorders. Rev Neurosci 2011, 22(2):153–169.

98. Steffan JS, Kazantsev A, Spasic-Boskovic O, Greenwald M, Zhu YZ, Gohler H, Wanker EE, Bates GP, Housman DE, Thompson LM: The Huntington’s disease protein interacts with p53 and CREB-binding protein and represses transcription. Proc Natl Acad Sci U S A 2000, 97(12):6763–6768.

99. Moily NS, Ormsby AR, Stojilovic A, Ramdzan YM, Diesch J, Hannan RD, Zajac MS, Hannan AJ, Oshlack A, Hatters DM: Transcriptional profiles for distinct aggregation states of mutant Huntingtin exon 1 protein unmask new Huntington’s disease pathways. Mol Cell Neurosci 2017, 83:103–112.

100. Chaturvedi RK, Hennessey T, Johri A, Tiwari SK, Mishra D, Agarwal S, Kim YS, Beal MF: Transducer of regulated CREB-binding proteins (TORCs) transcription and function is impaired in Huntington’s disease. Human Molecular Genetics 2012, 21(15):3474–3488.

101. Reiner A, Albin RL, Anderson KD, D’Amato CJ, Penney JB, Young AB: Differential loss of striatal projection neurons in Huntington disease. Proceedings of the National Academy of Sciences 1988, 85(15):5733–5737.

102. Sano H, Yasoshima Y, Matsushita N, Kaneko T, Kohno K, Pastan I, Kobayashi K: Conditional ablation of striatal neuronal types containing dopamine D2 receptor disturbs coordination of basal ganglia function. J Neurosci 2003, 23(27):9078–9088.

103. Kennedy L, Evans E, Chen CM, Craven L, Detloff PJ, Ennis M, Shelbourne PF: Dramatic tissue-specific mutation length increases are an early molecular event in Huntington disease pathogenesis. Hum Mol Genet 2003, 12(24):3359–3367.

104. Shelbourne PF, Keller-McGandy C, Bi WL, Yoon SR, Dubeau L, Veitch NJ, Vonsattel JP, Wexler NS, Group US-VCR, Arnheim N et al: Triplet repeat mutation length gains correlate with cell-type specific vulnerability in Huntington disease brain. Hum Mol Genet 2007, 16(10):1133–1142.

105. Gonitel R, Moffitt H, Sathasivam K, Woodman B, Detloff PJ, Faull RL, Bates GP: DNA instability in postmitotic neurons. Proc Natl Acad Sci U S A 2008, 105(9):3467–3472.

106. Swami M, Hendricks AE, Gillis T, Massood T, Mysore J, Myers RH, Wheeler VC: Somatic expansion of the Huntington’s disease CAG repeat in the brain is associated with an earlier age of disease onset. Hum Mol Genet 2009, 18(16):3039–3047.

107. Loupe JM, Pinto RM, Kim KH, Gillis T, Mysore JS, Andrew MA, Kovalenko M, Murtha R, Seong I, Gusella JF et al: Promotion of somatic CAG repeat expansion by Fan1 knock-out in Huntington’s disease knock-in mice is blocked by Mlh1 knock-out. Hum Mol Genet 2020, 29(18):3044–3053.

108. Kacher R, Lejeune FX, Noel S, Cazeneuve C, Brice A, Humbert S, Durr A: Propensity for somatic expansion increases over the course of life in Huntington disease. Elife 2021, 10.

109. Anderson CM, Swanson RA: Astrocyte glutamate transport: review of properties, regulation, and physiological functions. Glia 2000, 32(1):1–14.

110. Bak LK, Schousboe A, Waagepetersen HS: The glutamate/GABA-glutamine cycle: aspects of transport, neurotransmitter homeostasis and ammonia transfer. Journal of Neurochemistry 2006, 98(3):641–653.

111. Savchenko E, Teku GN, Boza-Serrano A, Russ K, Berns M, Deierborg T, Lamas NJ, Wichterle H, Rothstein J, Henderson CE et al: FGF family members differentially regulate maturation and proliferation of stem cell-derived astrocytes. Scientific Reports 2019, 9(1):9610.

112. Yang J, Luo X, Huang X, Ning Q, Xie M, Wang W: Ephrin-A3 reverse signaling regulates hippocampal neuronal damage and astrocytic glutamate transport after transient global ischemia. Journal of Neurochemistry 2014, 131(3):383–394.

113. Hassel B, Tessler S, Faull RL, Emson PC: Glutamate uptake is reduced in prefrontal cortex in Huntington’s disease. Neurochem Res 2008, 33(2):232–237.

114. Estrada-Sanchez AM, Montiel T, Segovia J, Massieu L: Glutamate toxicity in the striatum of the R6/2 Huntington’s disease transgenic mice is age-dependent and correlates with decreased levels of glutamate transporters. Neurobiol Dis 2009, 34(1):78–86.

115. Miller BR, Dorner JL, Shou M, Sari Y, Barton SJ, Sengelaub DR, Kennedy RT, Rebec GV: Up-regulation of GLT1 expression increases glutamate uptake and attenuates the Huntington’s disease phenotype in the R6/2 mouse. Neuroscience 2008, 153(1):329–337.

116. Sari Y, Prieto AL, Barton SJ, Miller BR, Rebec GV: Ceftriaxone-induced up-regulation of cortical and striatal GLT1 in the R6/2 model of Huntington’s disease. Journal of Biomedical Science 2010, 17(1):62.

117. Faideau M, Kim J, Cormier K, Gilmore R, Welch M, Auregan G, Dufour N, Guillermier M, Brouillet E, Hantraye P et al: In vivo expression of polyglutamine-expanded huntingtin by mouse striatal astrocytes impairs glutamate transport: a correlation with Huntington’s disease subjects. Hum Mol Genet 2010, 19(15):3053–3067.

118. Liévens JC, Woodman B, Mahal A, Spasic-Boscovic O, Samuel D, Kerkerian-Le Goff L, Bates GP: Impaired Glutamate Uptake in the R6 Huntington’s Disease Transgenic Mice. Neurobiology of Disease 2001, 8(5):807–821.

119. Shin JY, Fang ZH, Yu ZX, Wang CE, Li SH, Li XJ: Expression of mutant huntingtin in glial cells contributes to neuronal excitotoxicity. J Cell Biol 2005, 171(6):1001–1012.

120. Cross AJ, Slater P, Reynolds GP: Reduced high-affinity glutamate uptake sites in the brains of patients with Huntington’s disease. Neurosci Lett 1986, 67(2):198–202.

121. Hsu YT, Chang YG, Chern Y: Insights into GABA(A)ergic system alteration in Huntington’s disease. Open Biol 2018, 8(12).

122. Garret M, Du Z, Chazalon M, Cho YH, Baufreton J: Alteration of GABAergic neurotransmission in Huntington’s disease. CNS Neurosci Ther 2018, 24(4):292–300.

123. Santhakumar V, Jones RT, Mody I: Developmental regulation and neuroprotective effects of striatal tonic GABAA currents. Neuroscience 2010, 167(3):644–655.

124. Bayon-Cordero L, Ochoa-Bueno BI, Ruiz A, Ozalla M, Matute C, Sanchez-Gomez MV: GABA Receptor Agonists Protect From Excitotoxic Damage Induced by AMPA in Oligodendrocytes. Front Pharmacol 2022, 13:897056.

125. Bernstein EM, Quick MW: Regulation of gamma-aminobutyric acid (GABA) transporters by extracellular GABA. J Biol Chem 1999, 274(2):889–895.

126. Fenalti G, Law RH, Buckle AM, Langendorf C, Tuck K, Rosado CJ, Faux NG, Mahmood K, Hampe CS, Banga JP et al: GABA production by glutamic acid decarboxylase is regulated by a dynamic catalytic loop. Nat Struct Mol Biol 2007, 14(4):280–286.

127. Modi JP, Prentice H, Wu JY: Regulation of GABA Neurotransmission by Glutamic Acid Decarboxylase (GAD). Curr Pharm Des 2015, 21(34):4939–4942.

128. Wei J, Wu JY: Post-translational regulation of L-glutamic acid decarboxylase in the brain. Neurochem Res 2008, 33(8):1459–1465.

129. Kennedy L, Shelbourne PF, Dewar D: Alterations in dopamine and benzodiazepine receptor binding precede overt neuronal pathology in mice modelling early Huntington disease pathogenesis. Brain Res 2005, 1039(1-2):14–21.

130. Cepeda C, Starling AJ, Wu N, Nguyen OK, Uzgil B, Soda T, Andre VM, Ariano MA, Levine MS: Increased GABAergic function in mouse models of Huntington’s disease: reversal by BDNF. J Neurosci Res 2004, 78(6):855–867.

131. Rub U, Vonsattel JP, Heinsen H, Korf HW: The Neuropathology of Huntington s disease: classical findings, recent developments and correlation to functional neuroanatomy. Adv Anat Embryol Cell Biol 2015, 217:1–146.

132. Douaud G, Gaura V, Ribeiro MJ, Lethimonnier F, Maroy R, Verny C, Krystkowiak P, Damier P, Bachoud-Levi AC, Hantraye P et al: Distribution of grey matter atrophy in Huntington’s disease patients: a combined ROI-based and voxel-based morphometric study. Neuroimage 2006, 32(4):1562–1575.

133. Kassubek J, Bernhard Landwehrmeyer G, Ecker D, Juengling FD, Muche R, Schuller S, Weindl A, Peinemann A: Global cerebral atrophy in early stages of Huntington’s disease: quantitative MRI study. Neuroreport 2004, 15(2):363–365.

134. Beister A, Kraus P, Kuhn W, Dose M, Weindl A, Gerlach M: The N-methyl-D-aspartate antagonist memantine retards progression of Huntington’s disease. J Neural Transm Suppl 2004(68):117–122.

135. Ondo WG, Mejia NI, Hunter CB: A pilot study of the clinical efficacy and safety of memantine for Huntington’s disease. Parkinsonism Relat Disord 2007, 13(7):453–454.

136. Lucetti C, Del Dotto P, Gambaccini G, Dell’ Agnello G, Bernardini S, Rossi G, Murri L, Bonuccelli U: IV amantadine improves chorea in Huntington’s disease: an acute randomized, controlled study. Neurology 2003, 60(12):1995–1997.

137. Verhagen Metman L, Morris MJ, Farmer C, Gillespie M, Mosby K, Wuu J, Chase TN: Huntington’s disease: a randomized, controlled trial using the NMDA-antagonist amantadine. Neurology 2002, 59(5):694–699.

138. Galluzzi L, Vitale I, Aaronson SA, Abrams JM, Adam D, Agostinis P, Alnemri ES, Altucci L, Amelio I, Andrews DW et al: Molecular mechanisms of cell death: recommendations of the Nomenclature Committee on Cell Death 2018. Cell Death & Differentiation 2018, 25(3):486–541.

139. Mehta D, Jackson R, Paul G, Shi J, Sabbagh M: Why do trials for Alzheimer’s disease drugs keep failing? A discontinued drug perspective for 2010-2015. Expert Opin Investig Drugs 2017, 26(6):735–739.

140. Kim CK, Lee YR, Ong L, Gold M, Kalali A, Sarkar J: Alzheimer’s Disease: Key Insights from Two Decades of Clinical Trial Failures. Journal of Alzheimer’s Disease 2022, 87:83–100.

141. Hansson O, Petersen A, Leist M, Nicotera P, Castilho RF, Brundin P: Transgenic mice expressing a Huntington’s disease mutation are resistant to quinolinic acid-induced striatal excitotoxicity. Proc Natl Acad Sci U S A 1999, 96(15):8727–8732.

142. Hansson O, Castilho RF, Korhonen L, Lindholm D, Bates GP, Brundin P: Partial resistance to malonate-induced striatal cell death in transgenic mouse models of Huntington’s disease is dependent on age and CAG repeat length. J Neurochem 2001, 78(4):694–703.

143. Hansson O, Guatteo E, Mercuri NB, Bernardi G, Li XJ, Castilho RF, Brundin P: Resistance to NMDA toxicity correlates with appearance of nuclear inclusions, behavioural deficits and changes in calcium homeostasis in mice transgenic for exon 1 of the huntington gene. Eur J Neurosci 2001, 14(9):1492–1504.

144. Strong MK, Southwell AL, Yonan JM, Hayden MR, Macgregor GR, Thompson LM, Steward O: Age-Dependent Resistance to Excitotoxicity in Htt CAG140 Mice and the Effect of Strain Background. J Huntingtons Dis 2012, 1(2):221–241.

145. Edgar R, Domrachev M, Lash AE: Gene Expression Omnibus: NCBI gene expression and hybridization array data repository. Nucleic Acids Res 2002, 30(1):207–210.

