## Supplementary Figure 1 for "Evidence for glutamate excitotoxicity that occurs before the onset of striatal cell loss and motor symptoms in an ovine Huntington’s Disease model"

Supplementary 1 Multiplexed single nuclei RNA-seq libraries

| 10X Library | Samples | Case-control group |
| --- | --- | --- |
| 1 | HC373, HC337 | Control |
| 2 | HC382, HC335 | Control |
| 3 | HD372, HD317 | *OVT73* |
| 4 | HC357 | Control |
| 5 | HD377, HD339 | *OVT73* |
| 6 | HC334 | Control |
| 7 | HD376, HD383 | *OVT73* |
