## Supplementary Figure 2 for "Evidence for glutamate excitotoxicity that occurs before the onset of striatal cell loss and motor symptoms in an ovine Huntington’s Disease model"

| **10X library** | **1** | **2** | **3** | **4** | **5** | **6** | **7** |
| --- | --- | --- | --- | --- | --- | --- | --- |
| **Estimated #Nuclei** | 4,274 | 4,798 | 4,446 | 4,114 | 5,548 | 3,277 | 8,815 |
| **% Sequencing Saturation^1^** | 55.2 | 49.8 | 57.0 | 45.2 | 36.4 | 50.0 | 47.0 |
| **Total Reads** | 269,984,512 | 247,433,323 | 257,211,090 | 145,901,637 | 241,316,556 | 163,670,979 | 377,815,731 |
| **Total Genes detected** | 20,866 | 20,675 | 20,554 | 20,094 | 20,784 | 20,240 | 21,520 |
| **Median UMI/Barcode** | 1,902 | 1,766 | 2,162 | 2,064 | 2,386 | 2,212 | 2,612 |
| **Mean Reads/Barcode** | 63,169 | 51,570 | 57,852 | 35,465 | 43,496 | 49,945 | 42,861 |
| **Median Genes/Barcode** | 1,115 | 1,032 | 1,188 | 1,090 | 1,202 | 1,192 | 1,331 |
| **% Fraction Reads in Barcode** | 77.8 | 75.1 | 78.8 | 82.6 | 84.6 | 85.9 | 90.3 |
| **% Reads Mapped to Genome** | 98.9 | 98.7 | 98.9 | 98.9 | 98.6 | 98.9 | 96.0 |
| **% Reads Mapped to Transcriptome** | 25.4 | 30.6 | 32.8 | 32.5 | 32.6 | 33.1 | 40.0 |
| **% Reads Mapped to Exonic Regions** | 7.6 | 8.9 | 8.6 | 7.2 | 6.5 | 7.3 | 6.5 |
| **% Reads Mapped to Intronic Regions** | 57.8 | 55.1 | 55.7 | 56.9 | 57.3 | 56.3 | 53.4 |
| **% Reads Mapped to Intergenic Regions** | 27.6 | 28.4 | 28.9 | 29.8 | 29.5 | 30.4 | 30.2 |
| **% Reads Mapped Antisense to Gene** | 39.8 | 33.3 | 31.4 | 31.6 | 31.1 | 30.4 | 19.8 |
| **% Q30 Bases in Barcode** | 97.2 | 97.3 | 96.9 | 97.1 | 96.6 | 97.0 | 97.0 |
| **% Q30 Bases in RNA Read** | 91.6 | 91.7 | 90.6 | 91.4 | 91.0 | 91.5 | 91.3 |
| **% Q30 Bases in UMI** | 97.3 | 97.4 | 96.9 | 97.1 | 96.6 | 97.1 | 96.9 |

Supplementary 2 Summary statistics for multiplexed single nuclei RNA libraries

^1^Sequencing saturation defined as the fraction of reads originating from an already-observed UMI
