## Supplementary figures and images for "Evidence for glutamate excitotoxicity that occurs before the onset of striatal cell loss and motor symptoms in an ovine Huntington’s Disease model"

### Supplementary Figure 3

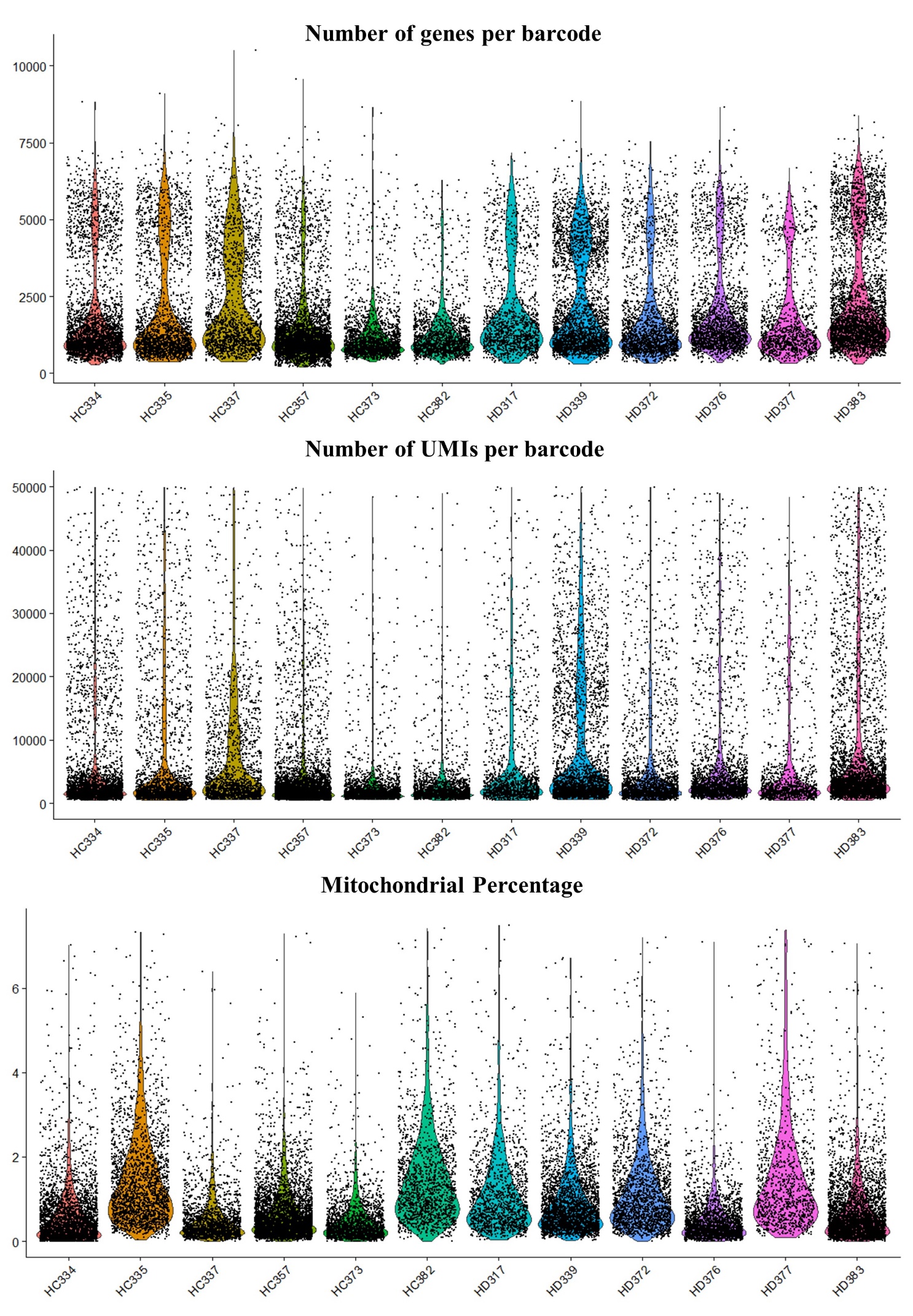
**Supplementary 3 Quality control metrics for single nuclei RNA libraries.**
