## Supplementary Figure 4 for "Evidence for glutamate excitotoxicity that occurs before the onset of striatal cell loss and motor symptoms in an ovine Huntington’s Disease model"

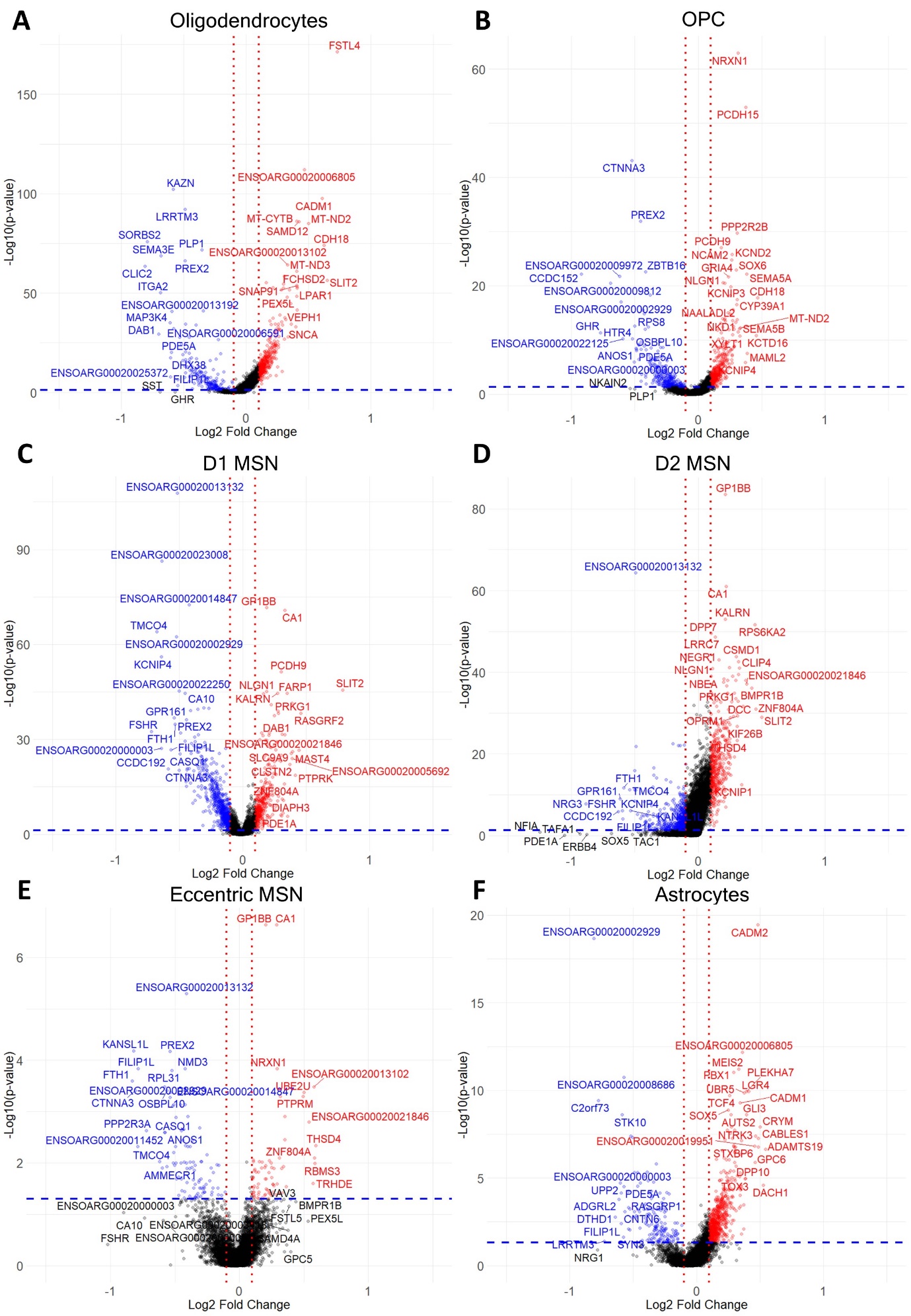

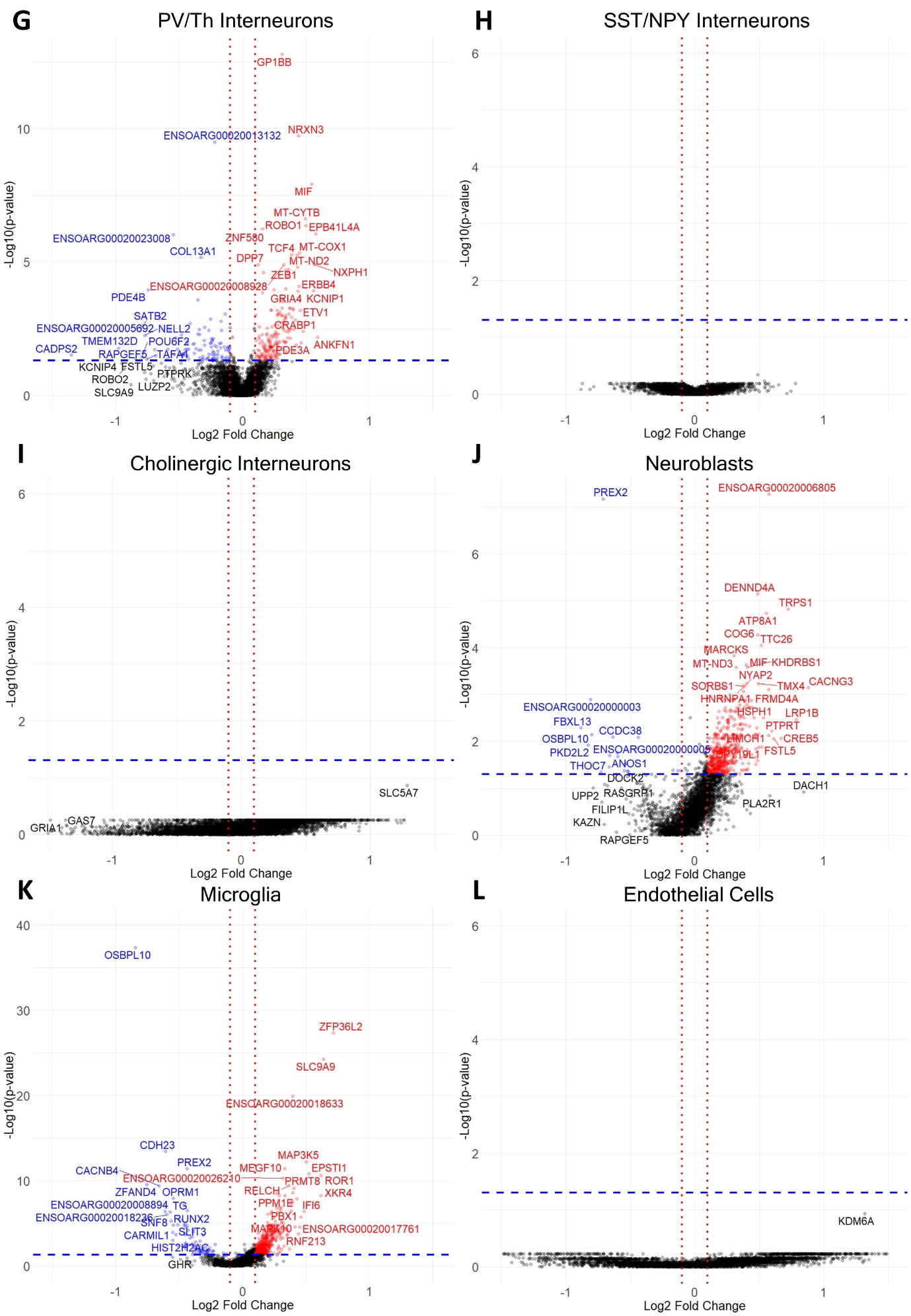

**Supplementary 4 Volcano plot of differentially expressed genes (DEGs) between *OVT73* and control for each cell type identified in the sheep striatum.** Horizontal blue line shown at p=0.05, vertical red lines shown at log2 fold change of -0.1 and 0.1.
