## Supplementary Figure 5 for "Evidence for glutamate excitotoxicity that occurs before the onset of striatal cell loss and motor symptoms in an ovine Huntington’s Disease model"

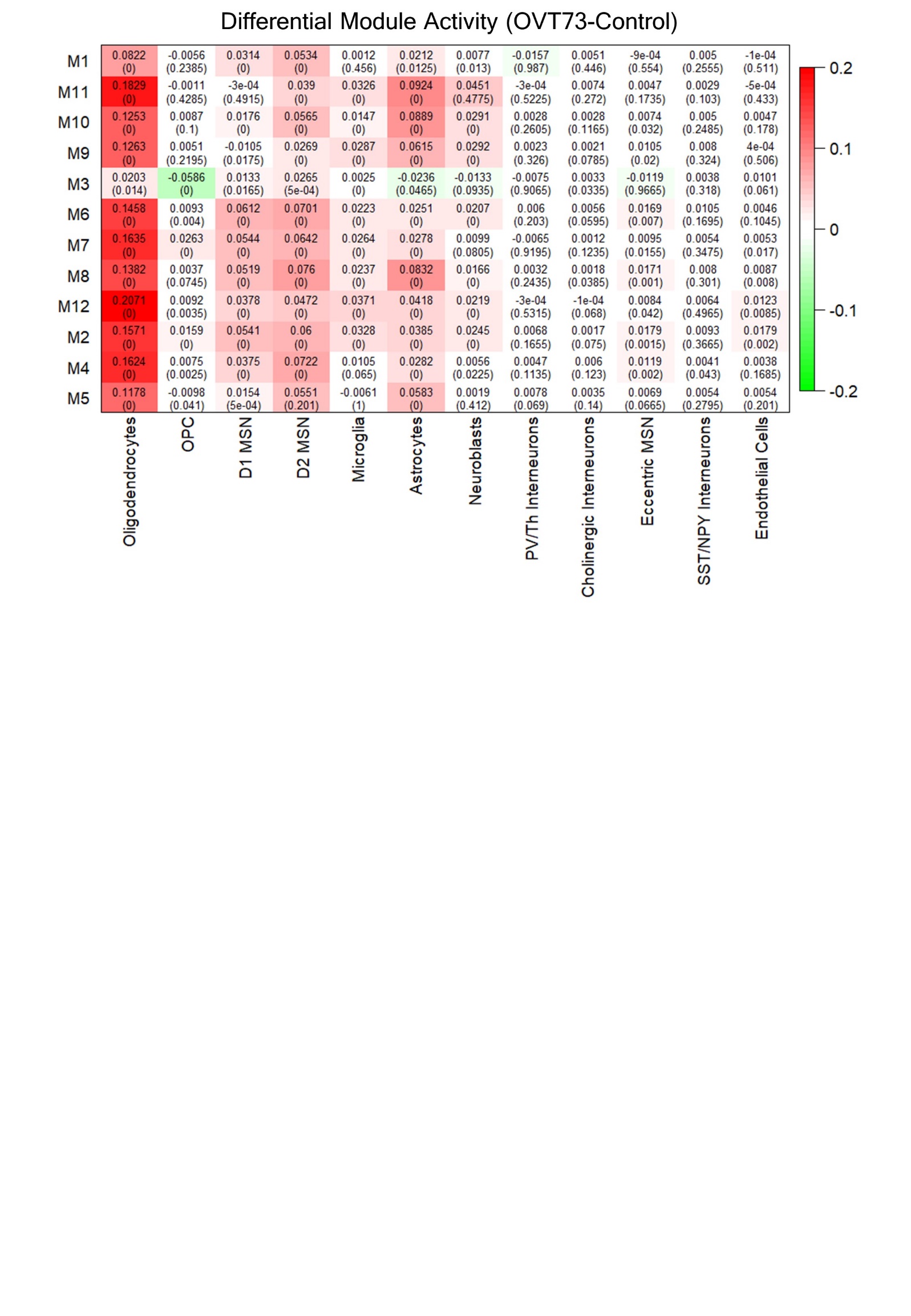

Supplementary 5 Differential co-expression module activity between *OVT73* and control cell types. Module activity in cell types were determined by computing the module eigengene (first principal component) using normalised expression values of module genes. Differential module activity was computed by subtraction of module eigengene values in *OVT73* and control cell types. A randomised permutation test with 2000 permutations was performed to determine significant differential module activity between *OVT73* and control cell types. P-values of the randomised permutation test are shown in the parentheses.
