## Supplementary Figure 6 for "Evidence for glutamate excitotoxicity that occurs before the onset of striatal cell loss and motor symptoms in an ovine Huntington’s Disease model"

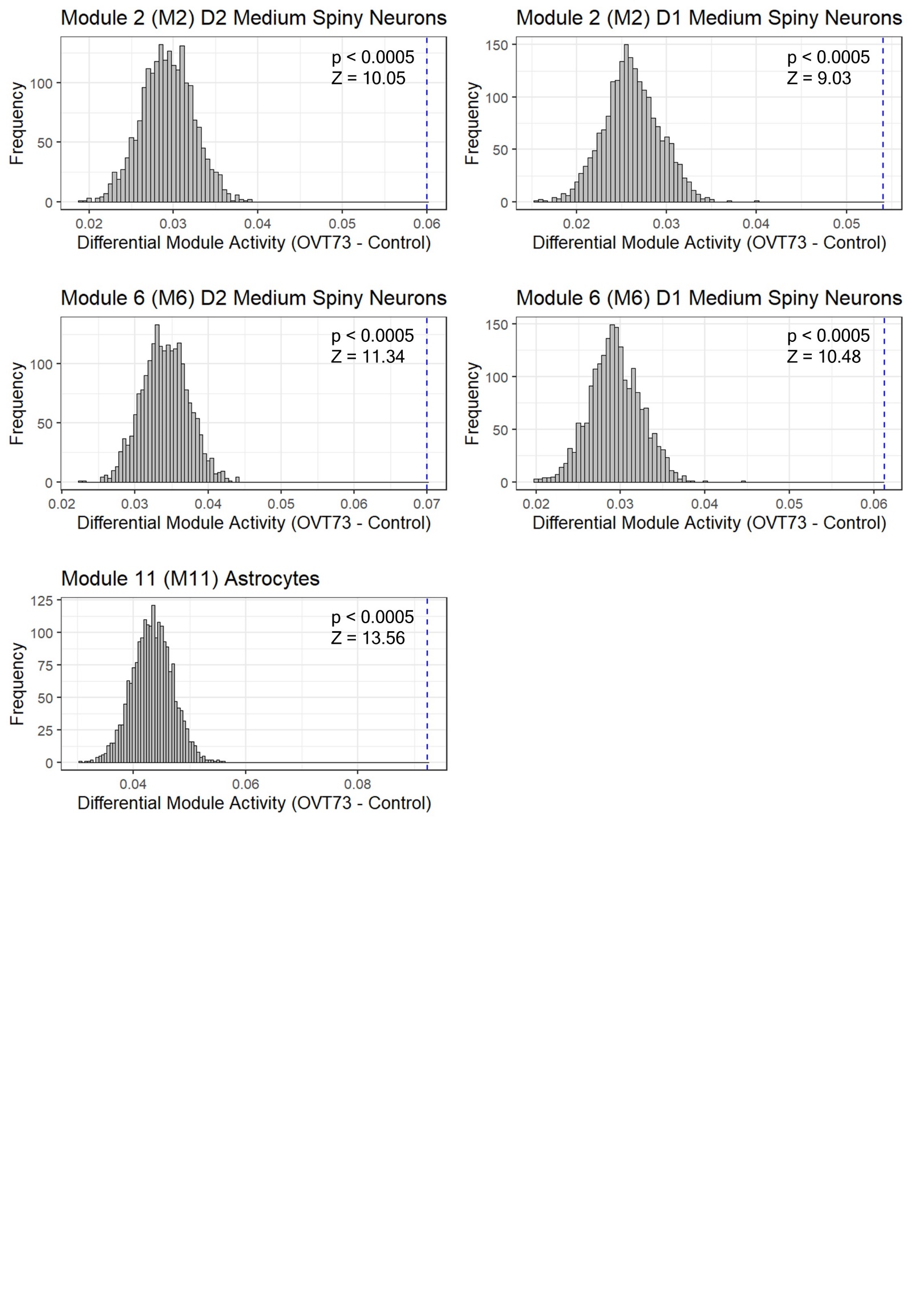
**Supplementary 6 Distribution of expected differential module activity scores between *OVT73* and control when genotype labels are randomly assigned.** Randomised permutation tests was performed with 2,000 permutations. Blue vertical line indicates actual differential module activity score.
