## Supplementary Figure 7 for "Evidence for glutamate excitotoxicity that occurs before the onset of striatal cell loss and motor symptoms in an ovine Huntington’s Disease model"

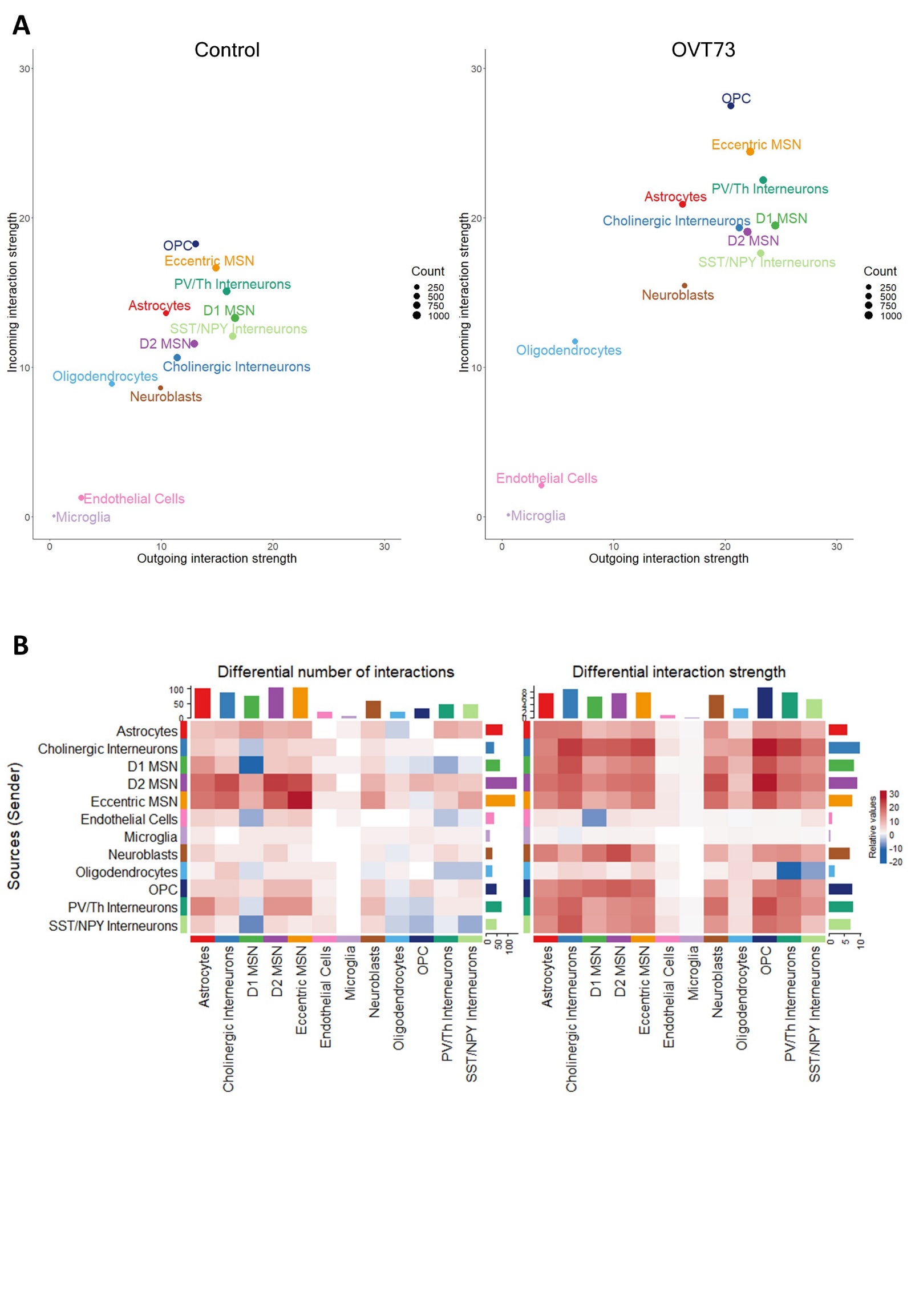

**Supplementary 7 Further visualisations of CellChat cell-cell communication networks.** (A) Comparison of the cell types that exhibit different degrees of incoming interaction signalling (from other cell types) and outgoing interactions signalling (to other cell types) in *OVT73* and control. An overall decrease in signalling was observed for the control dataset compared to *OVT73*. (B) Heatmap of differential number of interactions or differential interaction strength (communication probability) between any two cell types. Red represents increased signalling in the *OVT73* cell types compared to control, blue represents decreased signalling in the *OVT73* cell types compared to control. An overall increase in the number of interactions and interaction strength was observed for the *OVT73* cell types compared to control. The top-coloured bar plot represents the sum of the columns of values displayed in the heatmap (incoming signalling). The right bar plot represents the sum of row of values (outgoing signalling).
