## Supplementary Figure 8 for "Evidence for glutamate excitotoxicity that occurs before the onset of striatal cell loss and motor symptoms in an ovine Huntington’s Disease model"

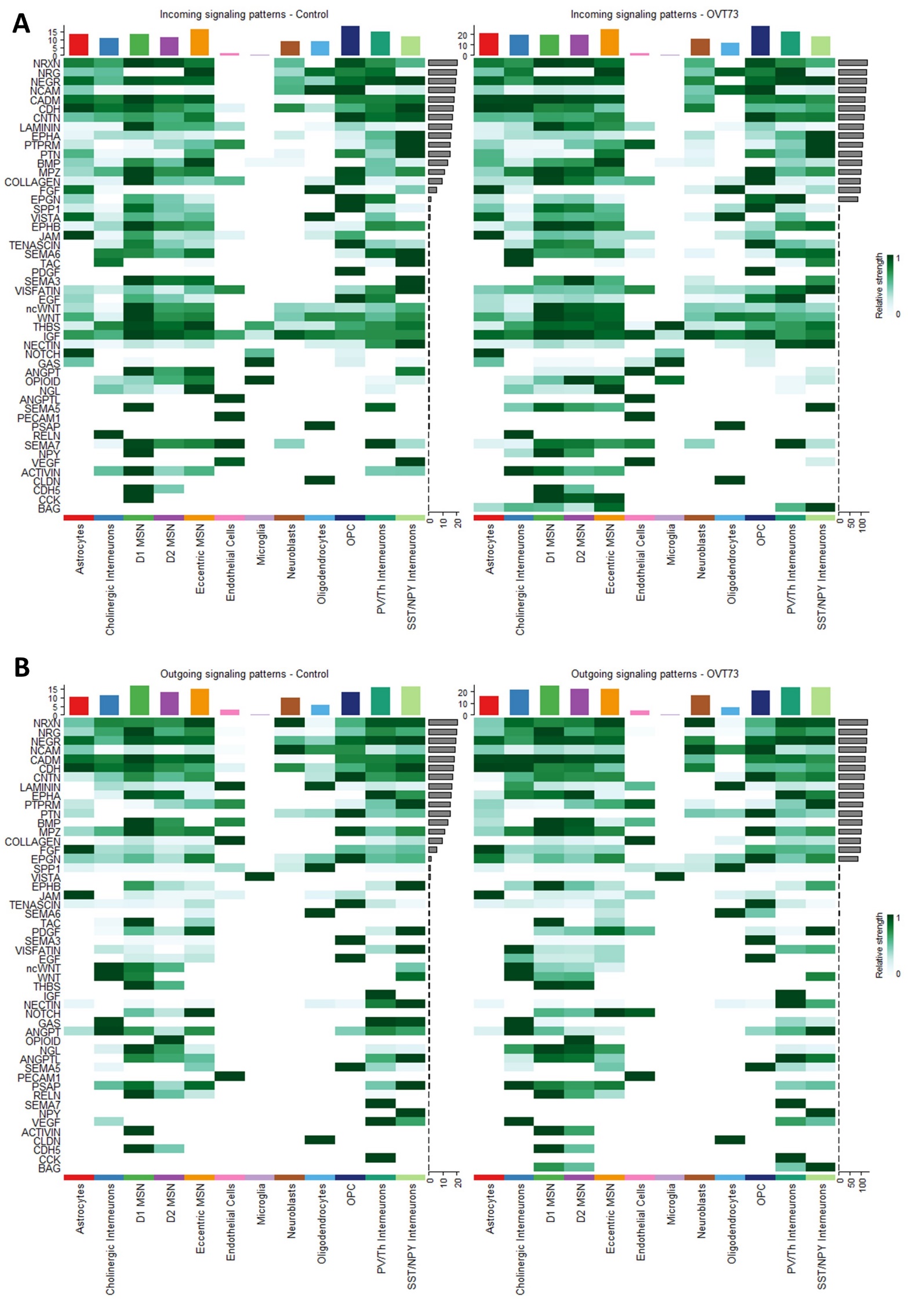
**Supplementary 8 Information flow from outgoing and incoming signalling pathways.** Information flow is defined as the sum of communication probabilities (strength) of all ligand receptor pairs in the signalling pathway.
