## Supplementary Figure 9 for "Evidence for glutamate excitotoxicity that occurs before the onset of striatal cell loss and motor symptoms in an ovine Huntington’s Disease model"

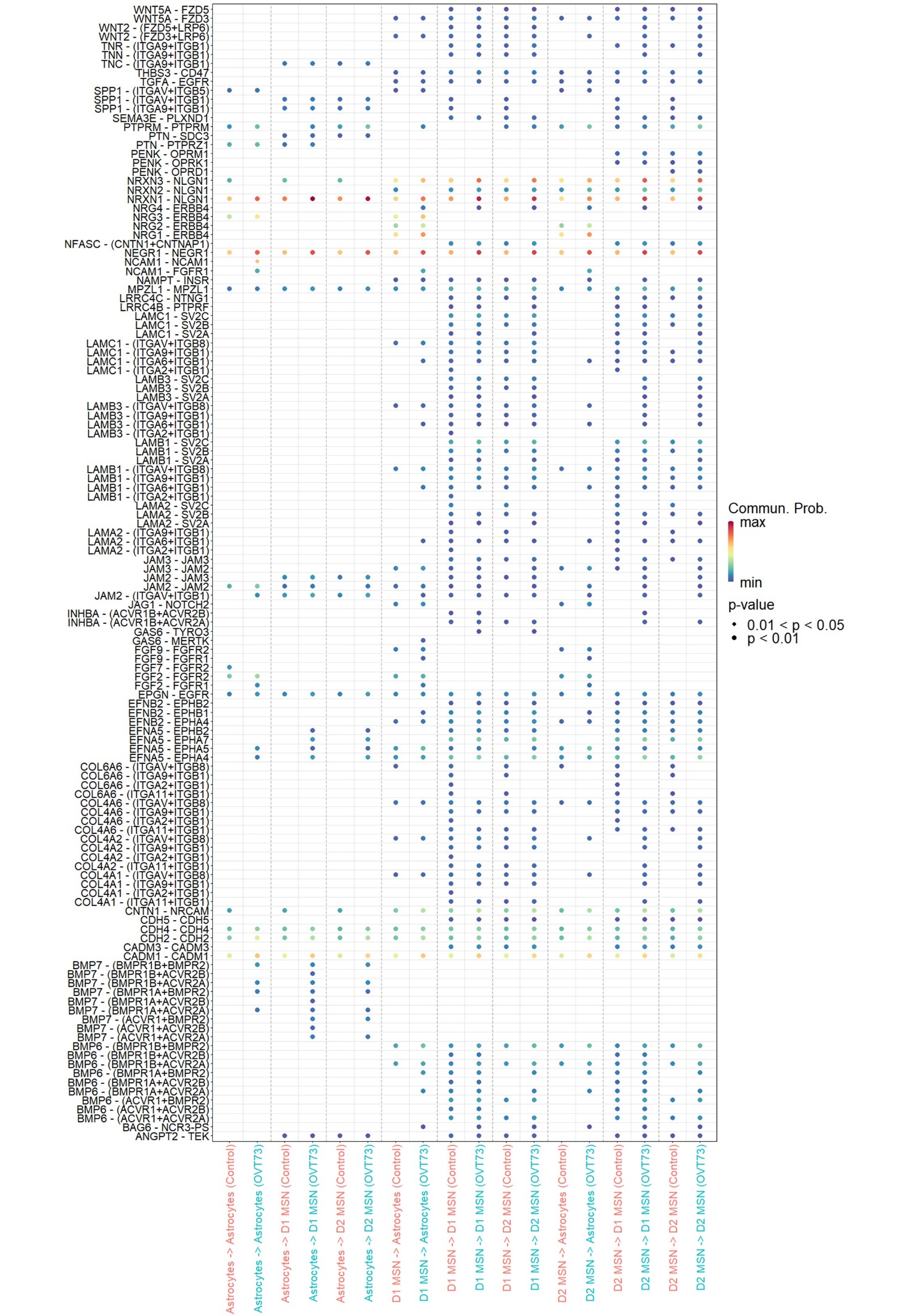
**Supplementary 9 Dot plot of ligand-receptor interactions for *OVT73* and control astrocytes and medium spiny neurons (D1, D2).** Color of the dot represents the communication probability; size of the dot represents the p value. Cell-cell communication probabilities were inferred from ligand receptor expression in single nuclei RNA-seq data using CellChat.
