## Supplementary Figure 10 for "Evidence for glutamate excitotoxicity that occurs before the onset of striatal cell loss and motor symptoms in an ovine Huntington’s Disease model"

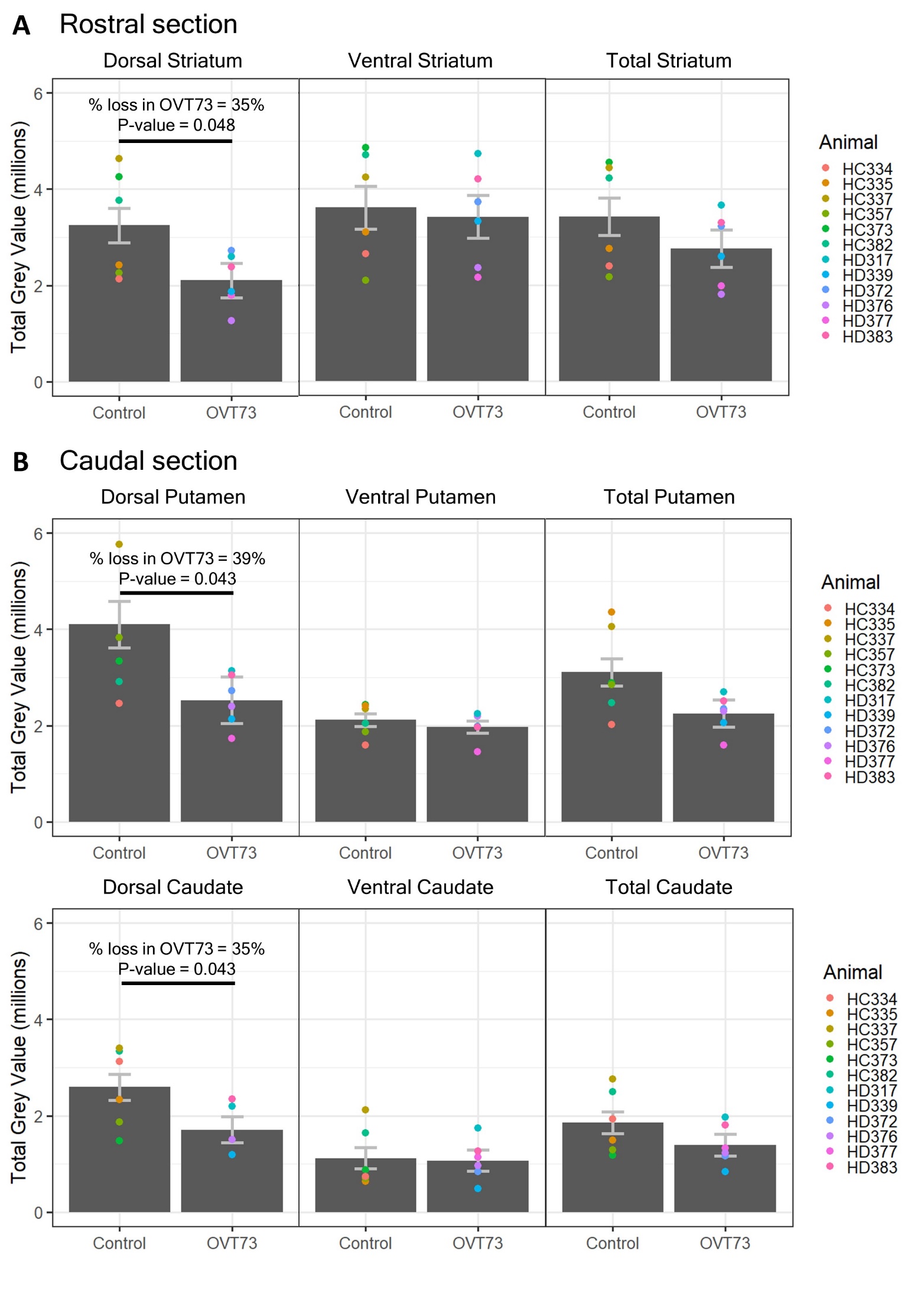

**Supplementary 10 Reduction in GABA_A_α1 immunoreactivity in the dorsal caudate, dorsal putamen and dorsal striatum of *OVT73* animals**. GABA_A_α1 immunoreactivity was reduced in the (A) dorsal striatum of the rostral section (35% reduction, p-value = 0.048) and (B) dorsal caudate (35% reduction, p-value = 0.043) and dorsal putamen (39% reduction, p-value = 0.043) of the caudal section of *OVT73* animals. The ventral striatum, ventral caudate and ventral putamen of rostral and caudal sections did not showcase a difference in GABA_A_α1 immunoreactivity.
