## Supplementary Figure 11 for "Evidence for glutamate excitotoxicity that occurs before the onset of striatal cell loss and motor symptoms in an ovine Huntington’s Disease model"

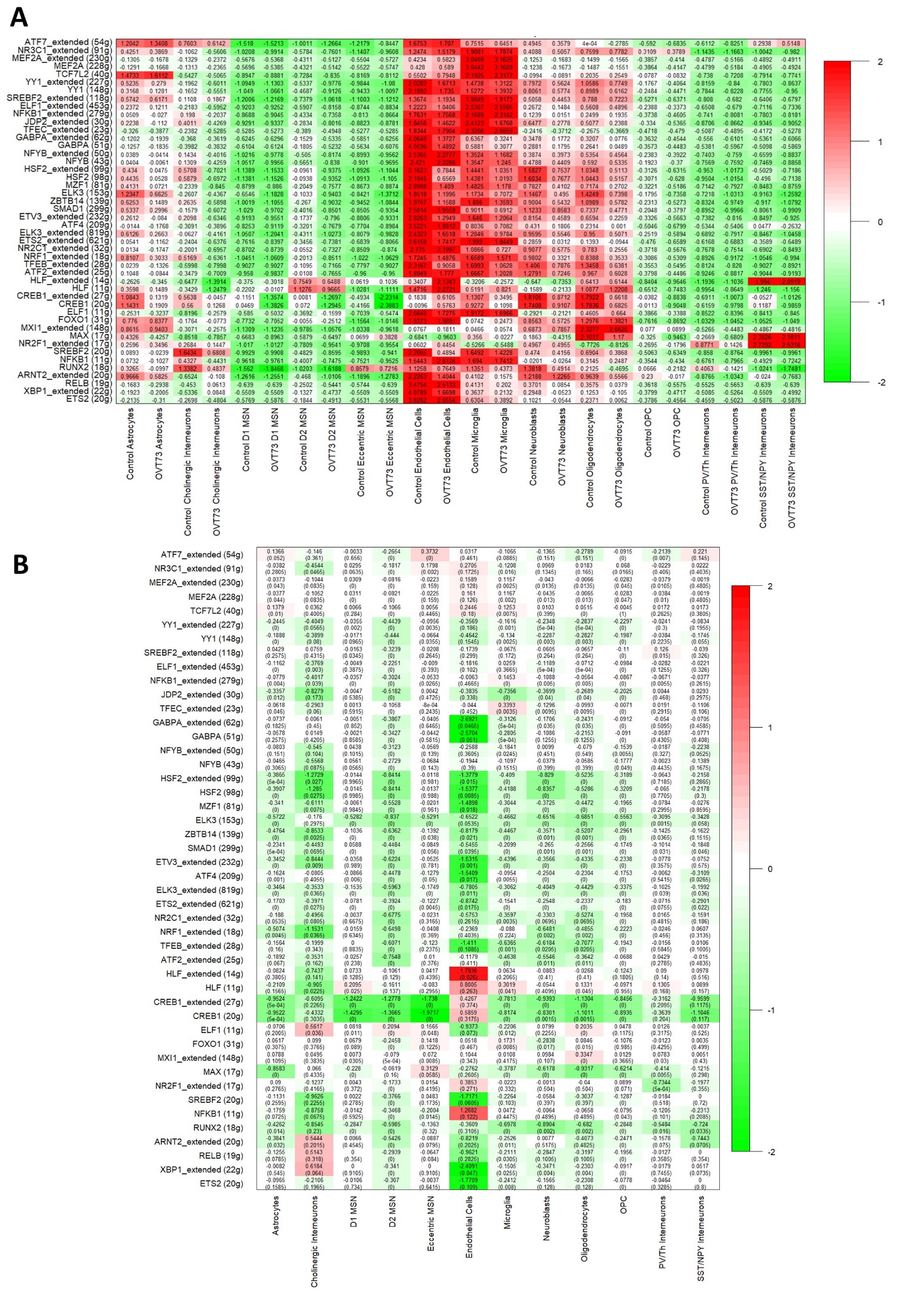
**Supplementary 11 Heatmap of gene regulatory network (regulon) activity in OVT73 and control cell types.** (A) Regulon activity was computed based on transcription factor regulated gene modules using OVT73 differentially expressed genes as input. A high regulon activity indicates genes within the regulon are positively regulated by the transcription factor. (B) Differential regulon activity was computed by subtraction of regulon activity in OVT73 and control cell types. A randomised permutation test with 2000 permutations was performed to determine significant differential regulon activity between OVT73 and control cell types. P-values of the randomised permutation test are shown in the parentheses.
