## Supplementary Figure 12 for "Evidence for glutamate excitotoxicity that occurs before the onset of striatal cell loss and motor symptoms in an ovine Huntington’s Disease model"

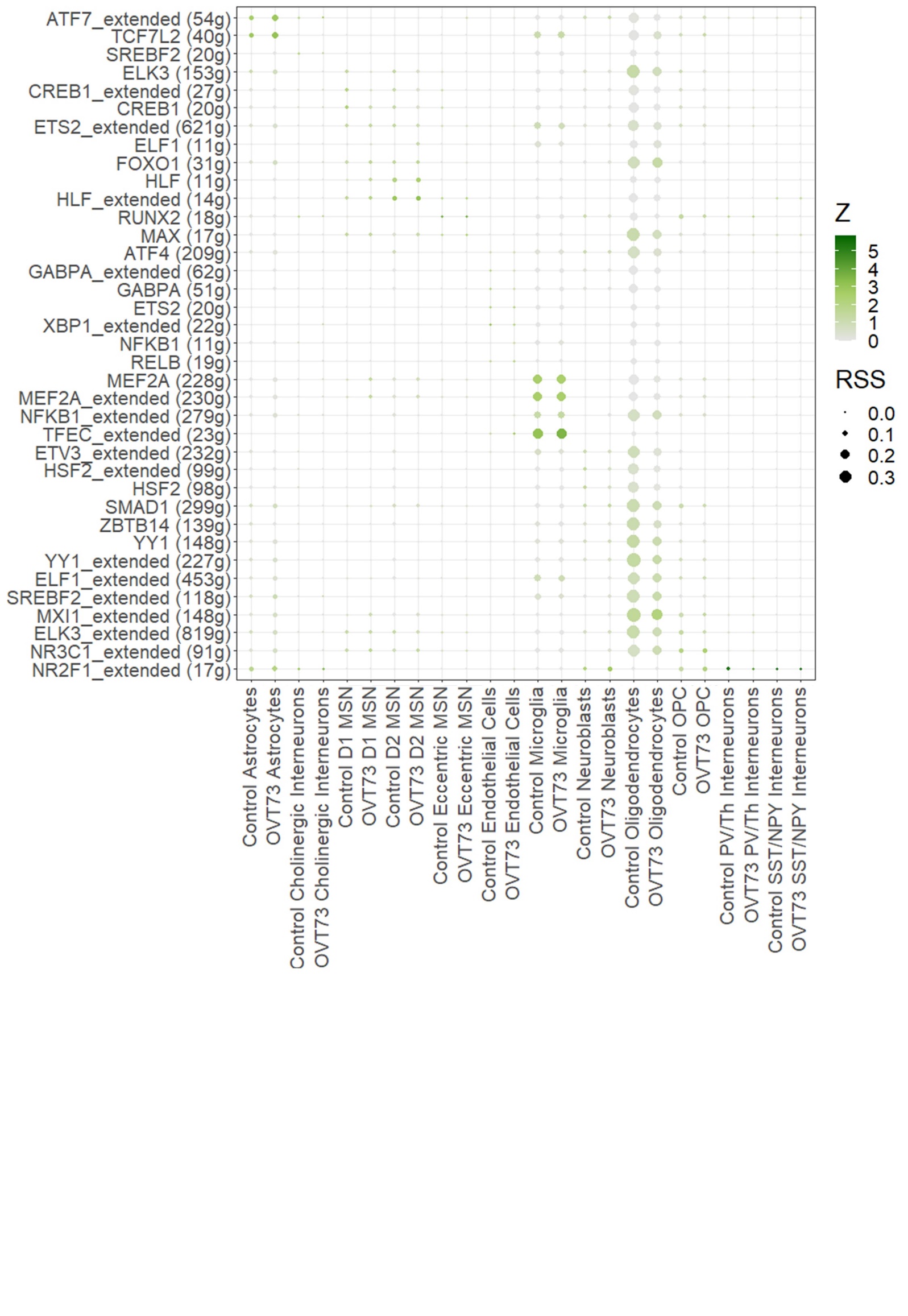

**Supplementary 12 Regulon specificity score of all regulons in the *OVT73* and control cell types.** The regulon specificity score assesses the exclusive regulon activity in cell types. Regulons in cell types with a specificity score of 1 indicates exclusive expression of the regulon in that one cell type, while a specificity score of 0 indicates the regulon is evenly expressed across all cell types. Size of the dot is proportional to the Z score.
