## Supplementary Figure 13 for "Evidence for glutamate excitotoxicity that occurs before the onset of striatal cell loss and motor symptoms in an ovine Huntington’s Disease model"

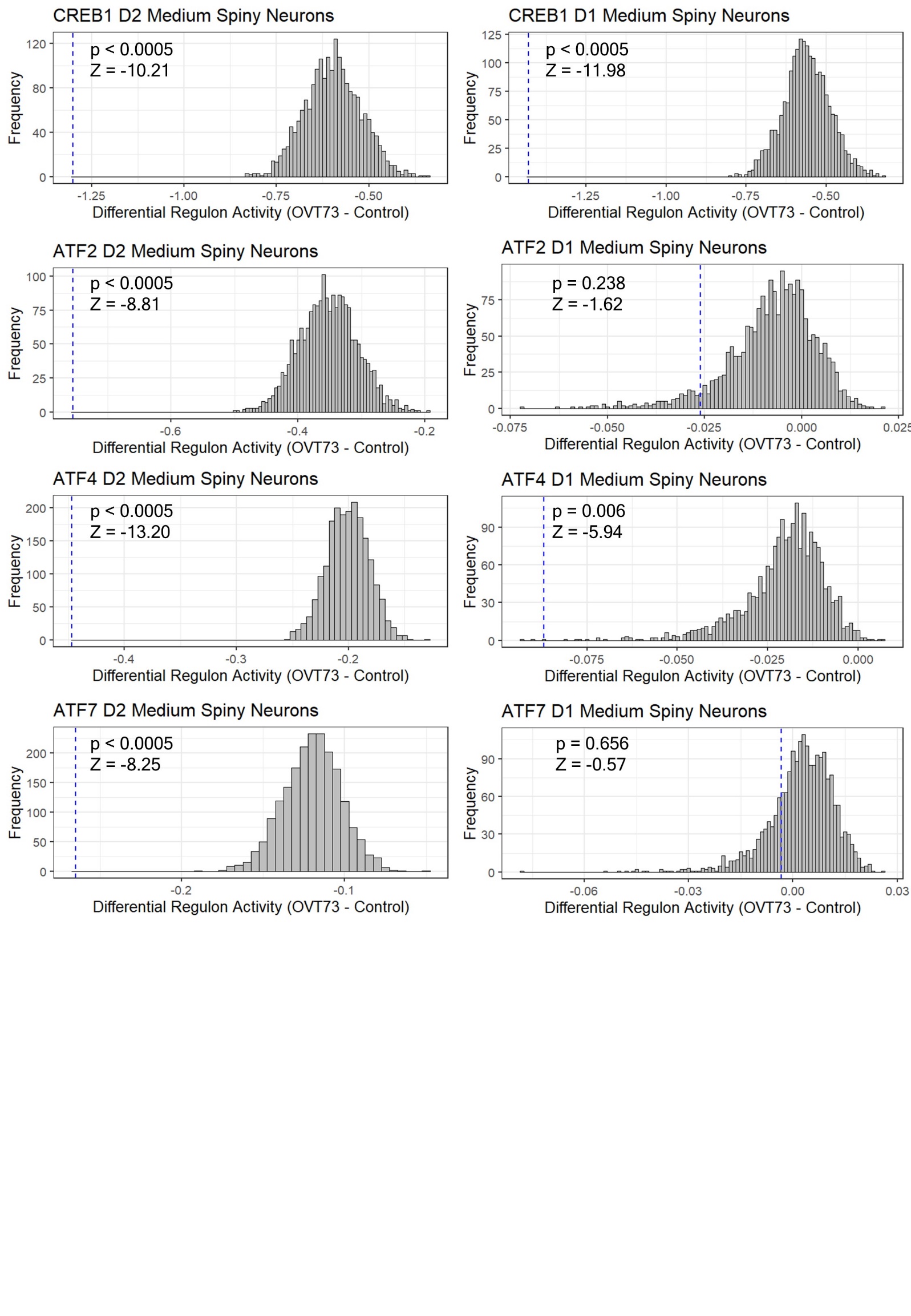

**Supplementary 13** **Distribution of expected differential regulon activity scores between *OVT73* and control when genotype labels are randomly assigned.** Randomised permutation tests were performed with 2,000 permutations. Blue vertical line indicates actual differential regulon activity score.
